# The Bioprocess TEA Calculator: An online techno-economic analysis tool to evaluate the commercial competitiveness of potential bioprocesses

**DOI:** 10.1101/2020.10.08.331272

**Authors:** Michael D. Lynch

**Affiliations:** Department of Biomedical Engineering, Duke University Durham, NC

## Abstract

Techno-economic analysis connects R&D, engineering, and business. By linking process parameters to financial metrics, it allows researchers to understand the factors controlling the potential success of their technologies. In particular, metabolic and bioprocess engineering, as disciplines, are aimed at engineering cells to synthesize products with an ultimate goal of commercial deployment. As a result it is critical to be able to understand the potential impact of strain engineering strategies and lab scale results on commercial potential. To date, while numerous techno-economic models have been developed for a wide variety of bioprocesses, they have either required process engineering expertise to adapt and/or use or do not directly connect financial outcomes to potential strain engineering results. Despite the clear value of techno-economic analysis, these challenges have made it inaccessible to many researchers. I have developed this online calculator (https://bioprocesstea.com OR http://bioprocess-tea-calculator.herokuapp.com/) to make the basic capabilities of early-stage techno-economic analysis of bioprocesses readily accessible. The tool, currently focused on aerobic fermentation processes, can be used to understand the impact of fermentation level metrics on the commercial potential of a bioprocess for the production of a wide variety of organic molecules. Using the calculator, I review the commercially relevant targets for an aerobic bioprocess for the production of diethyl malonate.

**Highlights:**

1. A generalized techno-economic analysis tool for aerobic fermentation based bioprocesses.
2. Relates strain and process improvements to projected commercial financial outcomes
3. Commercial bioprocess targets to produce diethyl malonate are estimated to include fermentation titers >150 g/L, rates > 5g/L-hr and > 90% yield, with downstream purification yields > 90% and purification costs limited to less than 20% of total.

## 1. Introduction

Biotechnology based fermentation processes have been successfully developed to produce a wide variety of molecules, from biologics and small molecule therapeutics to specialty, bulk and commodity chemicals, and even next generation biofuels. ^1–4^. Advances in fermentation science, synthetic biology, and metabolic and enzyme engineering are continually enabling novel or improved bioprocesses for a wide variety of these molecules. ^5–7^ At their core these efforts are focused on the development of technologies that may eventually lead to commercial products. Unfortunately, only a very small fraction of the amazing technical advances demonstrated over the past 30 years of metabolic engineering efforts have had a commercial impact. Additionally, many technologies that have moved toward commercialization have not reached success, relegating many exciting efforts to “proof of concept” studies. A critical challenge in the field is the ability to translate proof of concept lab efforts to commercial scale projects. There are many hurdles in the way of commercial success, ranging from a lack of funding, strain engineering pitfalls, and molecule specific technical challenges, to a lack of process robustness, reproducibility and scalability, ^8–10^ but one important generalized factor is routinely underestimating the technical performance required for commercial success.

One reason for underestimating commercially relevant performance metrics is a lack of routine techno-economic analysis (TEA) early in research programs. TEA is useful throughout the technology development lifecycle. When considering new ideas, pathways and molecules, innovators can use TEA to assess economic feasibility and potential. At the bench scale, scientists can use it to identify the process parameters that have the greatest effect on potential profitability. During process development, engineers can use TEA to compare the financial impact of different process conditions and configurations. TEA incorporates information from all of these stages of development, and provides a basis for making objective decisions. Furthermore, the time and cost needed for techno-economic analysis is small compared to that of running an experimental program. TEA is commonly used to assess the commercial potential of fermentation based processes for a wide variety of proteins and chemicals. Recent examples of molecule specific TEAs include those for terpenes, including limonene, adipic acid and other potential bulk or commodity chemicals as well as cellulosic biofuels. ^11–15^ However, to date many TEA tools are primarily based in complex spreadsheets or on complex software and not readily accessible by many researchers working in developing novel bioprocesses or biocatalysts. Broadly accessible TEA tools can benefit researchers from all backgrounds, working at all stages of development. Toward this goal, I have developed a freely available web based TEA application, the Bioprocess TEA Calculator (www.bioprocesstea.com, http://bioprocess-tea-calculator.herokuapp.com/). This tool is appropriate in the preliminary techno-economic assessment of conceptual bioprocesses that utilize aerobic fermentation and in understanding the potential impact of strain engineering efforts and fermentation performance metrics (including rate, titer and yield) on potential commercial feasibility. The calculator accepts a wide range of chemical targets and allows for significant user input (while also providing recommended defaults). I review the methodology underlying the calculator, as well as walk through a test use case: the identification of target performance metrics for a hypothetical aerobic bioprocess for the production of the chemical intermediate diethyl malonate.

## 2. Calculator Development

For bioprocesses, a plant can be divided into upstream and downstream areas as well as process utilities. Upstream areas are related to fermentation, while downstream areas are related to product purification. Process utilities are required and can be estimated for both upstream and downstream areas. Any TEA usually begins with, and is specific to, the details of the process being modeled, which historically would importantly include the target molecule being produced, the bioprocess performance, and the details of the downstream purification. This complexity can make a generalized TEA tool difficult to implement. As a result, I first sought to develop a generalized, yet meaningful, bioprocess model to enable reasonable cost estimates that are useful early in the strain and process development cycle. The approach taken is illustrated in Figure 1. Briefly, the Bioprocess TEA Calculator first accounts for the conversion of glucose to the product of interest (Figure 1a), calculating and accounting for theoretical conversion stoichiometry. Subsequently, (Figure 1b) an aerobic fermentation is modeled, wherein biomass growth is coupled to biosynthesis of the target chemical, given user defined performance metrics. This fermentation model is used for a detailed estimate of the operating (OPEX) and capital (CAPEX) costs of the bioprocess (Figure 1c), subject to user defined targets for plant performance (plant capacity for example) and downstream purification. As downstream purification (DSP) methods depend on the physical and chemical properties of the product, and other factors, like purity requirements and the fermentation byproduct profile, the current calculator only performs detailed estimates for primary cell removal, with remaining DSP costs estimated as a percentage of overall process costs. These DSP estimates can be altered by the user, either informed by a conceptual DSP process or by more general heuristics as discussed below. Lastly, estimates of OPEX and CAPEX are used along with financial inputs to perform a discounted cash flow analysis (Figure 1d) leading to estimated commercial outcomes, including the minimum selling price needed to achieve a target margin as well the net present value (NPV), and internal rate of return (IRR) for the project, based on the user defined target selling price.

**Figure 1:**
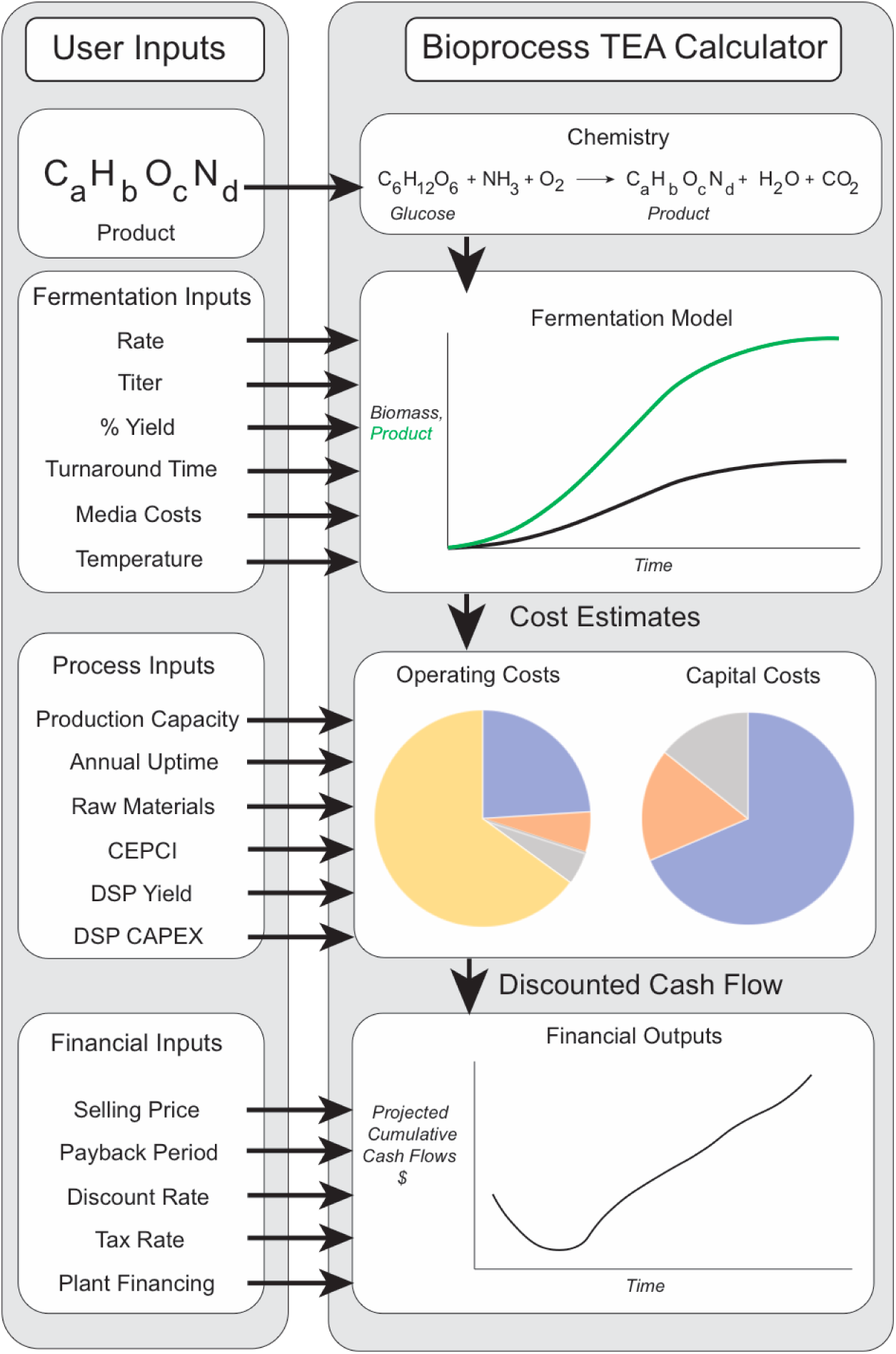
An overview of the Bioprocess TEA Calculator. The calculator is a responsive web application taking 4 primary types of user input : 1) the target chemical formula, 2) target bioprocess metrics and assumptions including rate titer and percentage of theoretical yield, media costs and temperature, 3) additional process inputs including production capacity, annual uptime, raw material cost, and assumptions on downstream purification (DSP) yield and costs, and 4) financial inputs including a target selling price, tax rate discount rate, and plant financing details. The calculator updates a techno-economic model in response to changes in these inputs and provides 4 key outputs. These include, 1) conversion stoichiometry, 2) a model of the fermentation, 3) operating and capital cost estimates, and 4) financial outputs based on a discounted cash flow analysis.

### 2.1 Chemistry

As illustrated in Figure 1, the first input into the calculator is the target chemical product. Currently, the calculator accepts molecules with carbon, hydrogen, oxygen and nitrogen atoms. While many algorithms can be used to balance chemical equations, ^16^ I chose to deploy a metabolically informed approach. Firstly, the problem is viewed from the perspective of the basic chemical building blocks used for biosynthesis, namely CO_2_ and H_2_. Glucose can be thought of as a combination of these two building blocks and specifically glucose provides 2 H_2_equivalents per carbon (CO_2_). Similarly, any potential organic chemical can be broken into carbon (CO_2_) and reducing equivalents (H_2_). Unless a chemical has the same H_2_/CO_2_ balance as glucose (an H_2_/CO_2_ ratio of 2) either carbon (CO_2_) or reducing equivalents (H_2_) will be the limiting reagent and so there are fundamentally three classes of “CHO” containing chemicals when considering glucose as a substrate. A chemical is either more reduced than glucose (an H_2_/CO_2_ ratio >2, for example ethanol or butanol), neutral with respect to glucose (an H_2_/CO_2_ ratio = 2, for example lactate), or more oxidized than glucose (an H_2_/CO_2_ ratio < 2, for example acetate or malonate). As Equations 1-6 illustrate, each of these cases leads to a different chemical equation that needs to be balanced. Algorithmically, the conversion of a given chemical input to H_2_ and CO_2_ is first considered, and the H_2_/CO_2_ ratio calculated, after which the appropriate final conversion equation is balanced. All of these equations can be balanced using simple algebra as outlined in Supplemental Materials Section 1. In the case of chemical products containing nitrogen a similar approach is used, except that all nitrogen atoms are assumed to originate from ammonia (NH_3_) which also brings reducing equivalents. This is consistent with basic biochemistry.

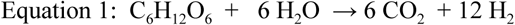

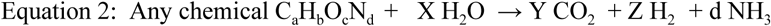

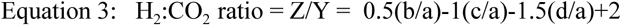

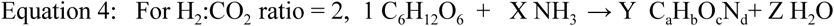

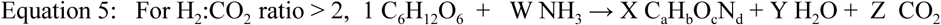

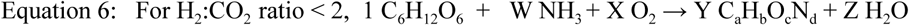

These calculations enable an estimate of the theoretical yield for the production of a given chemical from glucose. This calculation is based on the potential theoretical chemical conversion and does not assume a particular biosynthetic pathway or account for thermodynamics. Using a particular biochemical pathway may result in a theoretical pathway yield lower than the theoretical chemical yield. Additionally, while certain conversions may be stoichiometrically possible they may not be energetically feasible. Additionally, yield coefficients for ammonia and oxygen, used in calculations in the fermentation model below, are calculated at this step. These yield coefficients are essentially the grams of product per gram of ammonia (or oxygen). Due to the details of the stoichiometric calculations, the current version of the calculator assumes glucose (dextrose) as the carbon and electron source for the production of any given target chemical. Future improvements may include additional feedstocks.

### 2.2 Fermentation Model

With the theoretical chemistry defined, I developed a basic model for aerobic fermentation. This model takes into account user defined inputs such as the target titer, volumetric production rate and percentage of theoretical yield as illustrated in Figure 1. For more details regarding the fermentation model, please refer to Supplemental Materials Section 2. In this model the percentage of theoretical yield is based on the theoretical chemical yield described above. The user should account for differences in yield attributable to a lower theoretical pathway yield for any specific biochemical pathway. For simplicity, the calculator assumes growth associated production and logistic growth. The target yield along with the target titer define the final amount of product and dictate the available sugar (not used for product synthesis) which can be used for biomass growth, as described in Figure 2. Biomass is modeled as a byproduct chemical (C_3.85_H_6.69_O_1.78_N) with a molecular weight of 95.37 g/mole, which is produced from glucose at 80% of theoretical yield. ^17,18^. As a result the user input titer and percent of theoretical yield are used to estimate the final biomass levels at the end of fermentation.

**Figure 2:**
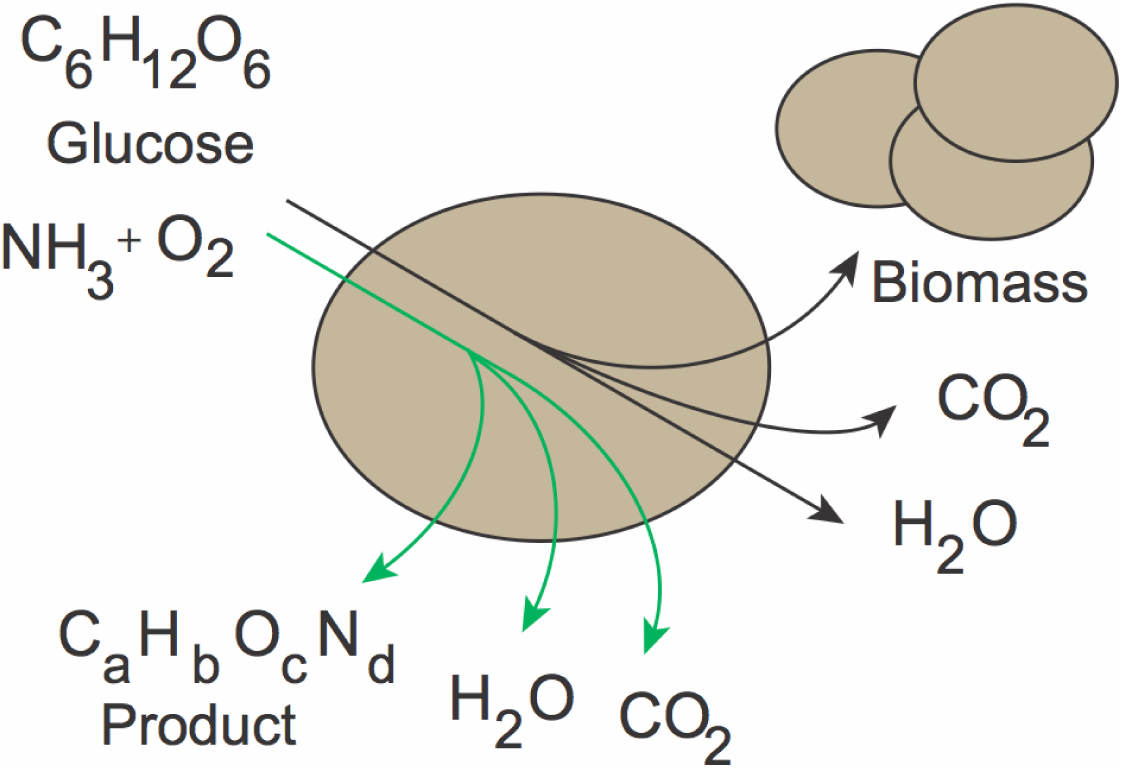
A depiction of growth associated product biosynthesis as modeled via the Bioprocess TEA Calculator. Glucose, ammonia, and oxygen are the sole feedstocks used for biomass (black line) or product (green line) synthesis. The theoretical yield and stoichiometry of product biosynthesis (green pathway) is first calculated based on the product formula, giving a theoretical yield. The user defines the percentage of this theoretical yield to be achieved in the fermentation. The product conversion assumes 100% of theoretical yield for the glucose distributed to the product. This percentage (capped at 99%) defines the available sugar remaining that may be used for biomass growth (back lines). The biomass conversion assumes 80% of theoretical yield with 20% lost to the complete oxidation of glucose. This distribution is used to calculate the product to cell ratio or amount of product that is produced per gram of dry cell weight produced. This model, together with the user defined production rates and yield, is used to estimate final biomass levels and growth rates.

The total fermentation time is calculated from the titer and average volumetric production rate. The starting biomass level is assumed to be 1% of the final biomass (based on a 1% inoculum of a similarly dense seed culture). As mentioned a logistic growth rate is assumed and calculated based on final and initial biomass levels and the fermentation time. Finally the product to cell ratio, a measure of growth associated product biosynthesis, is calculated. This ratio is in effect the gram of product made per gram of biomass as the cells grow. This can be used to estimate a logistic product formation rate.

Together the above calculations lead to a fermentation time course, which can be used for basic material and energy balances and to estimate process requirements including raw materials, oxygen demand and oxygen transfer rates and cooling requirements, which in turn can be used with other inputs to estimate operating and capital costs.

### 2.3 Downstream Purification

Currently, the calculator specifically estimates the costs associated with primary cell removal, which is modeled as centrifugation. These costs are standard in most bioprocesses, whether the product is in the supernatant or the cell pellet. However, the calculator currently only enables the user to estimate additional downstream purification operating and capital costs as a fraction of total costs without any detailed estimates of these unit operations. Importantly, the current estimates assume a single byproduct: carbon dioxide. All dextrose not going to produce biomass or product is completely oxidized to carbon dioxide. In most cases this likely overestimates the oxygen requirements of the process and represents a worse case scenario with respect to aeration, however this also represents a best case scenario with respect to DSP. Real world downstream purification processes must often deal with the removal of several unwanted organic byproducts, which are a function of the product and biosynthetic pathway, as well as strain and process conditions deployed.

Capital costs associated with DSP can range widely depending on the complexity of the DSP and number of unit operations required. Simple distillation, such as is used for ethanol, may be on the lower end (∼ 20%), whereas DSPs requiring chromatographic separations or electrodialysis would be anticipated to account for 50-70% of total capital costs. ^14,19–21^ Operating costs for DSP are often a smaller fraction of the total percentage of OPEX, and I recommend from 20% at the low end to 50% at the high end. ^22^ Figure 3, adapted from data presented by Straathof ^22^ relates the reported percent of production costs attributed to downstream purification for a variety of molecules. These data can be used for rough estimates of DSP costs for different types of products.

**Figure 3:**
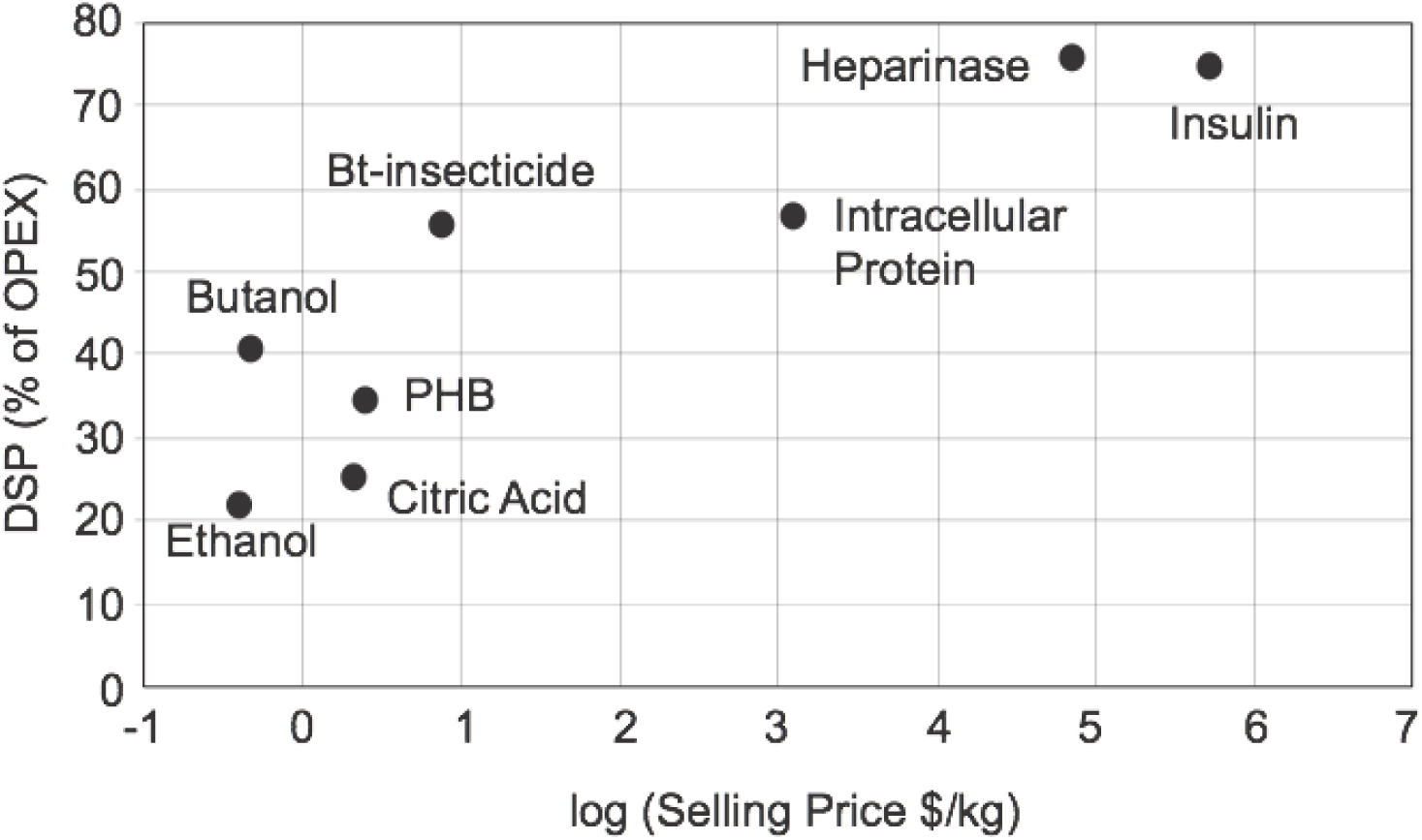
The contribution (%) of downstream purification to production costs as a function of the log selling price for various reported commercial bioprocesses. Data are adapted from Straathof, 2011. ^22^ Abbreviations: PHB - polyhydroxybutyrate, Bt (*Bacillus thuringiensis*)-insecticide.

### 2.4 Operating and Capital Cost Estimates

The current version of the Bioprocess TEA Calculator assumes the construction of a new facility in a Build-Own-Operate (BOO) Financial model. Estimated operating and capital costs assume a general plant configuration as illustrated in Figure 4. Briefly the facility can be broken down into the following 6 key areas including 1) Main fermentation 2) Seed fermentation, 3) Primary cell removal, 4) Process utilities, 5) Downstream purification, and 6) associated infrastructure (warehousing and office buildings, site development etc). In this brownfield design utilities for the process are modeled but main gas lines and electrical supply are provided to the facility. Additionally, while waste is initially heated (to kill biomass) and pH adjusted, a wastewater treatment plant is not considered in this design, and would need to be provided via colocation.

**Figure 4:**
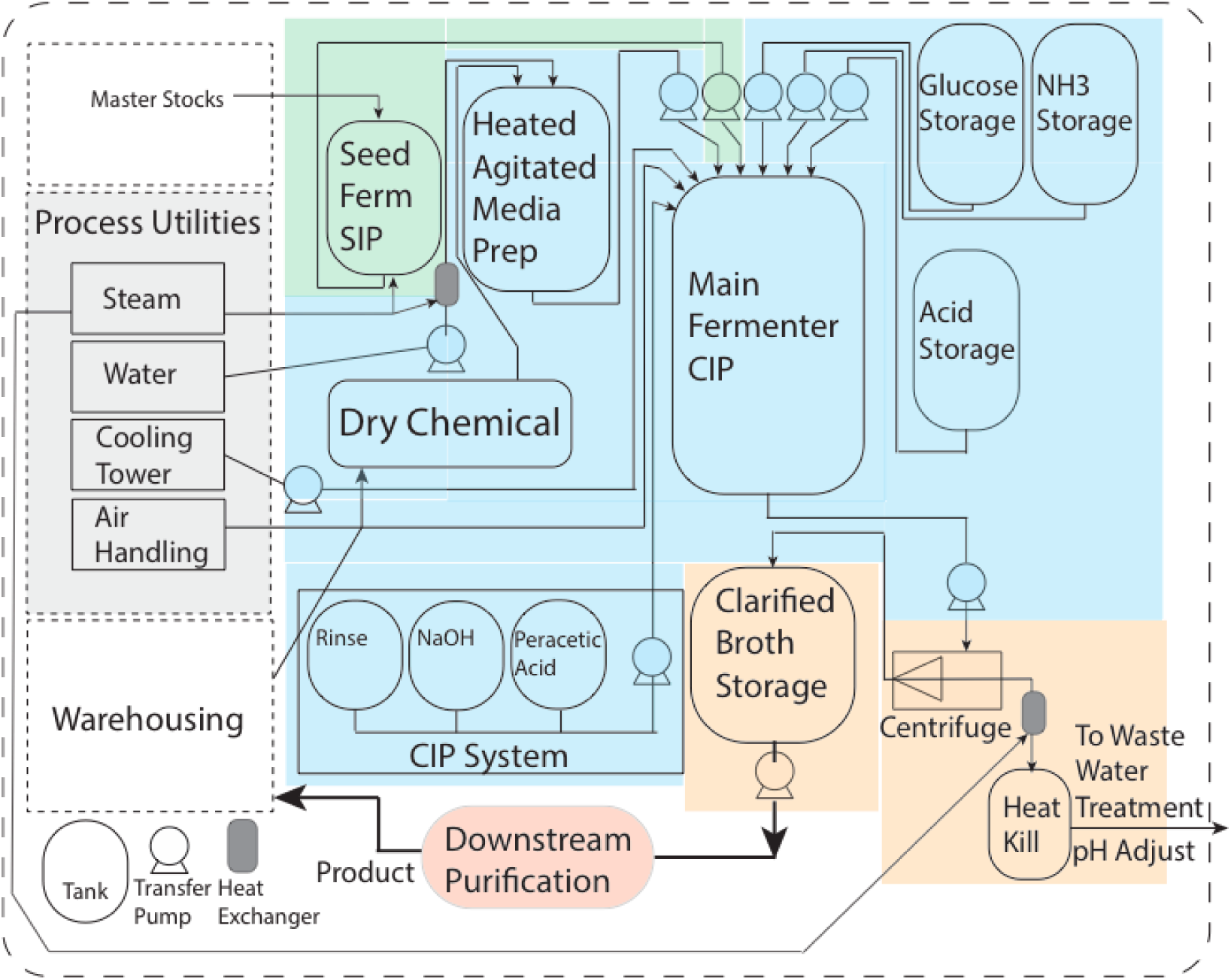
A diagram of the generalized bioprocessing facility as modelled by the Bioprocess TEA Calculator. The facility is divided into six key areas modeled to various degrees, the main fermentation (highlighted in blue), the seed fermentation train (highlighted in green), primary cell removal (highlighted in orange), process utilities (highlighted in gray), downstream purification (highlighted in red) and accessory facilities (offices, warehousing, etc.). Equipment costs and installation factors for tanks, transfer pumps, heat exchangers and centrifuges are calculated based on user defined inputs, as are specific equipment and installation costs associated with process utilities including steam generation, cooling, water and air handling. Piping costs are assumed to be 4.5% of total installed equipment costs (TIC). Additional site development including buildings and other costs are estimated as a percentage of the TIC. The boundary of the facility (dashed lines) does not include basic utility generation (electricity, natural gas) or final wastewater treatment.

Operating costs are estimated prior to capital costs, as many of the capital estimates depend on the OPEX. Operating costs estimates include variable costs such as raw materials and utilities as well as fixed operating costs including labor and additional fixed costs. For more details on specific calculations underlying estimates refer to Supplemental Materials Sections 3 and 4. The calculator estimates the raw materials and process utilities needed given the target plant capacity and fermentation performance as well as other user defined inputs. Raw materials and utilities estimated include: water, glucose, ammonia, and the costs of CIP (clean in place) reagents, electricity, steam and compressed air. Glucose costs are estimated based on the user defined glucose price as well as the overall fermentation yield and yield in the DSP. First the amount of glucose needed for product biosynthesis and biomass production are calculated as described above to give a target fermentation yield. The fermentation yield together with the overall DSP yield (as input by the user) defines the total glucose needed. The default glucose pricing is at $0.18/lb which corresponds to bulk 2020 pricing for DE95 corn syrup, which is usually > 650 g/L dextrose. Lower “over the fence” prices may be achievable when co-locating a facility with a sugar mill. Ammonia costs are estimated similarly to glucose costs. In this case the ammonia required for product synthesis is estimated from the stoichiometry, if the product contains nitrogen. Similarly the amount of ammonia required for biomass synthesis is estimated based on final biomass levels and the stoichiometry defining biomass production from glucose and ammonia. The default ammonia price of $0.12/lb reflects 2020 estimates.

The costs of media components are estimated based on the final biomass levels in the main production fermentations. As a result the user inputs the cost of media components per kg or biomass (kgCDW). The default value of ∼$0.40/kgCDW assumes a minimal mineral salts media, based on FGM25 minimal media (which can support biomass levels of 25gCDW/L) as reported by Menacho-Melgar et al and detailed in Supplemental Materials. ^23^ It is important to note that complex media, such as LB and derivatives, are very expensive and can easily cost in the range of $25-$100/kgCDW. The costs for the clean in place (CIP) of the main fermentation vessels is based on two cleaning solutions: a caustic solution (NaOH) and a peracetic acid sterilizing solution, for which cost estimates are detailed in the Supplemental Materials.

Cost estimates for the majority of the utilities in this calculator are based on utility estimations by Ulrich & Vasudevan ^24^. The estimates rely on the Chemical Engineering Plant Cost Index (CEPCI a metric of inflation, https://www.chemengonline.com/pci-home), and the “Cost of Fuel, ($/GJ)” which is in turn estimated based on the user defined natural gas cost.

Labor costs are estimated based on the number of main fermentation tanks needed to meet target plant capacity according to Davis et al. ^15^ Similarly, other fixed operating costs are estimated as a percentage (3.7%) of the direct capital investment (discussed below) which scales with the size of the plant. ^15^

Based on the fermentation model and other process inputs as well as estimated operating expenses calculated above, the size of major required equipment can be estimated. Specifically, the size and or number of tanks, transfer pumps, heat exchangers and other key items including centrifuges, tanks and pumps for water treatment and processing, boilers, cooling towers (and cooling tower pumps), and air handling (compressors and receivers) are all estimated. For details on these calculations refer to Supplemental Materials Section 4. Once needed equipment sizes are estimated, a standard approach to capital estimation is used, which relies on scaling factors and installation factors (as well as accounting for inflation) according to Equations 7 and 8 and detailed in Supplemental Materials, Section 4. ^15,25,26^ Additionally, plant piping is estimated at 4.5% of total installed equipment costs and the capital costs for the seed fermentation area are estimated at 27% of the main fermentation area, without more detailed estimates.^27^ Using the total of installed equipment costs in all key areas, the total capital investment is then estimated as outlined in Table 1 below.

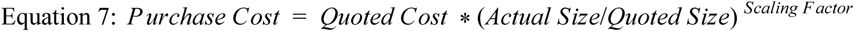

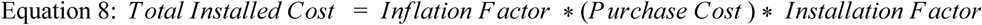

**Table 1:** Capital Cost Estimates ^28^

| **Table 1: Capital Cost Estimates** <sup>28</sup> |  |
| --- | --- |
| **Direct Costs** |  |
| Site Development | 0.09* Total Installed Equipment Costs |
| Warehousing | 0.04* Total Installed Equipment Costs |
| Administrative Buildings | 0.05*Total Installed Equipment Costs |
| Total Direct Costs | = Total Installed Equipment Costs +<br>Warehousing + Administrative Buildings +Site Development |
| **Indirect Costs** |  |
| Proratable Expenses | 0.1*Total Direct Costs |
| Field Expenses | 0.1*Total Direct Costs |
| Office & Construction Fees (Home Office) | 0.2*Total Direct Costs |
| Other Start Up Costs | 0.1*Total Direct Costs |
| Project Contingency | 0.1* Total Direct Costs |
| Total Indirect Costs = | Proratable Expenses + Field Expenses + Home Office + Project<br>Contingency + Other Start Up Costs |
| **Total Capital Expenses** |  |
| Fixed Capital Investment (FCI) | Total Direct Costs + Total Indirect Costs |
| Working Capital (WC) | 0.05*FCI |
| Total Capital Investment (TCI) | FCI +WC |

### 2.5 Financial Outputs

Estimated operating and capital costs can be used, with additional financial assumptions, to 1) calculate a minimum selling price (MSP) and 2) perform a discounted cash flow (DCF) analysis. Key financial inputs include assumptions on plant financing such as the fraction of the plant financed, loan term and interest rate. Additionally, the user defined target margin is used for calculating the MSP, and other user defined inputs including the target selling price, project payback period and discount rate, are used in the DCF analysis.

For the purposes of this calculator, the MSP is the selling price required to achieve the target (user defined) margin upon the completion of plant ramp up (ie. the first year the facility is producing at nameplate production capacity). The calculator defaults to a 2 year construction period, followed by a 3 year period needed to ramp to full production. The MSP calculation accounts for the target margin, operating expenses, depreciation and ongoing capital expenditures as well as loan payments (both principal and interest). Importantly, the MSP may be higher than a reasonable market price in which case the process needs optimization to be commercially competitive or is commercially infeasible.

The DCF analysis uses projected expenses and revenues along with the discount rate and payback period to estimate a net present value (NPV), return on investment (ROI) and internal rate of return (IRR) for the project. Discounted cash flow analysis is a method of evaluating the financial value of a project taking into account the time value of money. Specifically the “project” in this case, is the investment in constructing and starting a new production facility for the target product and the estimated returns based on projected future sales up until the payback period. For details on these financial calculations refer to Supplemental Materials Section 5. Key Definitions can be found in Supplemental Materials Section 0.

Importantly, neither the margin nor MSP, affect any calculations in the DCF analysis, which is based on the selling price. When using the calculator, the MSP gives you an estimate of the minimum price at which you need to sell the product to make your target margin. The DCF gives you estimates of the NPV given a specific payback period and discount rate and importantly a target selling price, which should be competitive in the market. The DCF also calculates an ROI and IRR which are dependent on the payback period but not the discount rate. I often recommend the IRR as the metric to choose when evaluating/comparing potential projects. IRR is the annual rate of growth an investment is expected to generate. IRR is calculated using the same concept as NPV, except it sets the NPV equal to zero. The calculator defaults to a minimum IRR of -99.99%.

### 2.6 Web Application Development

Finally, the Bioprocess TEA calculator web application was developed using JavaScript and Node.js. (https://nodejs.org) All calculations are performed client side (in the browser) and saved only when the user chooses to save an analysis for a more detailed download. Currently, all saved analyses are viewable by all users. MongoDB (https://www.mongodb.com/) is currently used as the database and the site upon initial launch was deployed through Heroku (www.heroku.com). Screen captures of key inputs and outputs are illustrated in Figure 5. Any TEA can be saved. Once saved, a detailed report can be downloaded including all major results and costs estimates including the fermentation time course, a Proforma (projected cashflows) and detailed equipment and capital costs.

**Figure 5:**
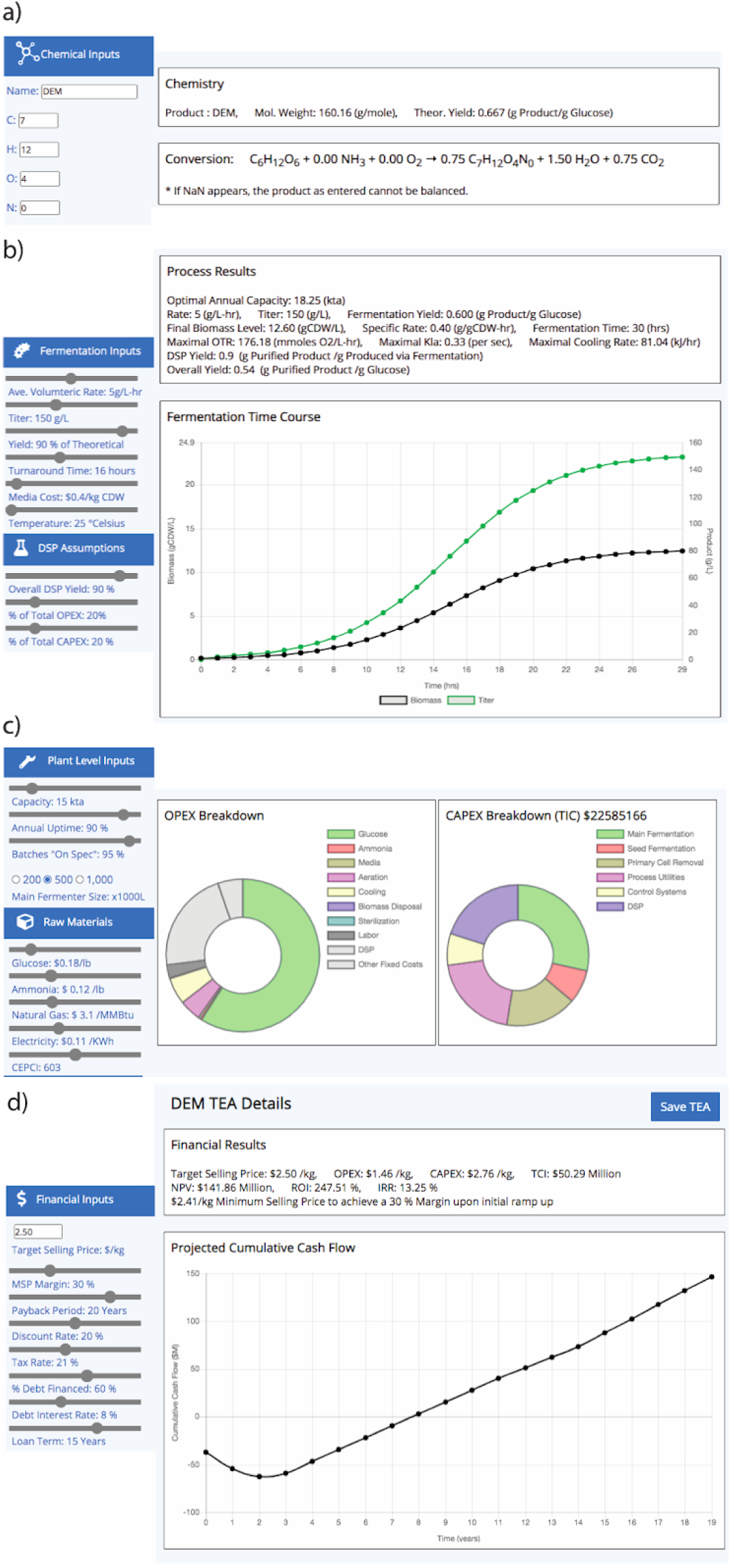
Screen captures of the Bioprocess TEA Calculator’s inputs and respective outputs. a) product inputs and chemistry outputs, b) fermentation and downstream purification performance inputs and fermentation outputs, c) additional plant and raw material inputs and estimated output costs and d) financial inputs and outputs. Users have the option to save a given TEA, and after saving export a csv file with all analysis details.

## 3. Use Cases and Limitations

As mentioned above, this calculator should be used for projects requiring the construction of new dedicated plants for the production of chemicals using bioprocesses and should be used for processes producing anywhere from 1 to 100kta of a chemical. This equates to 1,000,000 to 100,000,000 kg per year. Processes resulting in annual production volumes lower than this should not use these estimates. Additionally, for products at the low end of these production volumes, contract manufacturing (leveraging existing facilities, rather than building a new plant) may be commercially viable and a better path. The current version of this calculator assumes new construction. The user should assess whether the volumes considered by this calculator are meaningful in light of the market size for a given product. For example, if the current worldwide annual production of a given chemical is lower than 1 kta, this calculator is likely not appropriate. It is important to note that the current capital cost estimates are for brownfield development. This assumes that the plant is constructed on a site with current access to a wastewater treatment facility and basic utilities such as natural gas and electric power.

The calculator uses cost estimates that would be aligned with chemical or food grade production. Food grade production requires the ability to demonstrate cleanability of all process equipment as well as related quality control. The current estimates should not be used for pharmaceutical grade estimates. Pharmaceutical grade production would require a 15-30% increase in equipment costs, as specific equipment requirements often need to be certified by manufacturers, increasing prices. Additionally, pharmaceutical production requires significant quality control systems and expensive “sterile” areas not contemplated by the calculator. These add significant capital and operating costs (including labor) which would need to be considered.

As mentioned, DSP costs beyond primary cell removal (Figure 4) are rough estimates. The unit operations required for any purification are driven by the physiochemical properties of the product as well many other factors including target purity specifications and the fermentation byproduct profile. The simplest and lowest-cost chemical separations are typically vapor-liquid separations at low temperature and near-ambient pressure, such as distillation and stripping. As mentioned above, downstream processing for products in these categories may be as low as 20% of overall production costs. ^22^ More complicated DSP processes may incur much higher costs. Values for DSP of 20% of OPEX and 50% of CAPEX are recommended when the user does not have a specific downstream in mind, although I always recommend evaluating the impact of changing these cost estimates on the potential commercial feasibility of a given bioprocess.

Currently the calculator does not account for product acidity or basicity. Specifically it assumes the product will have a pKa ∼ 7.0, and that the pH of the fermentation is held constant at neutral pH. This means that for products which are acids and/or bases the model does not account for the costs of the titrants needed for neutralization, although capital estimates for titrant delivery are included. While future iterations of the calculator may include these estimates based on user input pKas, currently, if necessary, the user can estimate the worst case scenario. This can be done by assuming one mole of base or acid required per mole of product produced in the facility. Default ammonia prices of $0.12/lb ($0.000264/g) correlate to ∼ $/0.0045/mole of base. Currently, I would recommend a sulfuric acid price estimate of $150/tonne ($0.000150/g) which correlates to ∼ $/0.0074/mole of acid. In the event that the uncharged form of a product is desired after purification, the user can estimate a worse case scenario where an equivalent amount of both acid and base are needed in the process, for example 1 mole of base needed per mole of product in the fermentation to maintain a neutral pH and 1 mole of acid in the DSP to convert the salt form of the product to the neutral form. For example, in the case of a 50 kta facility producing acrylic acid (MW = 72.06g/mole), these calculations would add an additional annual titrant cost of $8,257,000 or $0.16/kg ($0.072/lb), which in the case of a commodity is not insignificant. Disposal of the large amount of salt waste in the DSP would also increase the relative DSP operating and capital costs. Of course the impact of titration costs are magnified in the example of acrylic acid and are likely less of an issue for most products.

## 4. A technoeconomic analysis of an aerobic bioprocess for diethyl malonate production

Lastly, I present a use case for the Bioprocess TEA Calculator, namely the production of diethyl malonate (DEM). DEM, is also an important intermediate in the chemical syntheses of vitamins B1 and B6, barbiturates, non-steroidal anti-inflammatory agents, other numerous pharmaceuticals including chloroquine, as well as agrochemicals.^29^ Additionally, malonic acid esters can also be functionalized to create reactive molecules including methylene malonates which can be used in advanced adhesives. ^30,31^ DEM is currently produced from petroleum sources at estimated annual volumes > 60 kta, in a non-sustainable and environmentally harmful process utilizing sodium cyanide. ^29^ Current estimated selling prices for DEM range from $2.50-$3.00/kg. ^29,32^ This monomer is a great candidate for renewable production from sugar feedstocks via fermentation, with a theoretical conversion as illustrated in Figure 6.

**Figure 6:**
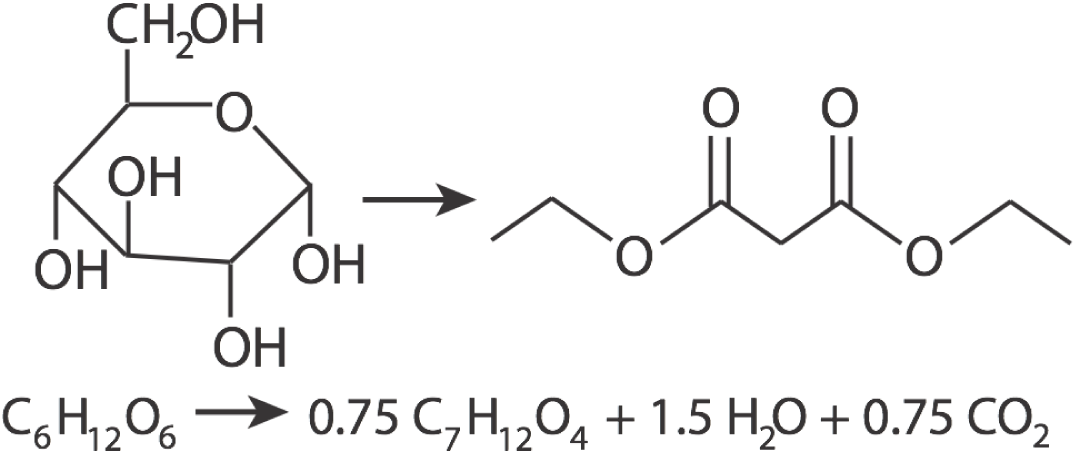
The conversion of glucose to diethyl malonate

After entering the formula for DEM in the calculator, as illustrated in Figure 5, the user can evaluate the impact of other key process inputs on the fermentation performance, cost breakdown and financial outputs. In this case, the calculator can be used systematically to estimate commercially relevant metrics for a DEM bioprocess. In this analysis, I set the DEM selling price to $2.50/kg to be competitive with petroleum based processes/ The plant capacity was set to 15 kta or one quarter of the current estimated market size. A fermentation yield of 90% of theoretical was assumed and held constant. DEM is volatile and so a DSP process reliant on fractional distillation with costs similar to that of ethanol purification were assumed, specifically DSP OPEX costs were set to 15% and DSP CAPEX costs to 20%. Other default assumptions were not initially varied and set as illustrated in Figure 5.

First, the impact of the key performance metrics including titer, rate and DSP yield was evaluated. Results are given in Figure 7a. A titer > 90g/L, DSP yield > 90% with an average volumetric rate > 0.9g/L-hr is required to achieve a positive IRR on the project, leading to an estimated MSP <$5.00/kg, and a total capital investment of < $80 Million USD. These metrics do not lead to a commercially attractive project and so further increases in volumetric rate were considered as shown in Figure 7b. A volumetric rate > 2.5 g/L is needed to achieve an MSP < $2.60 with an estimated TCI ∼ $60 Million USD. However even with these production rates, the estimated IRR is still less than 10%. As mentioned above IRR is often used as a metric of the financial potential of a project, and although a heuristic, a projected IRR of 15-20% is often considered a go/no-go when considering large capital projects. To further evaluate process optimization opportunities to increase the project IRR, increases in rate and titer were evaluated as illustrated in Figure 7c-e. A rate of 5g/L-hr and a titer of 150g/L is estimated to be required to achieve an IRR of ∼18%. These metrics result in an estimated TCI of ∼ $46 Million USD and a MSP ∼ $2.30/kg. Lastly, I evaluated the impact of the initial capital cost assumptions on these estimates as illustrated in Figure 8. In this analysis I held the fermentation performance metrics constant (Rate: 5g/L-hr, Titer: 150g/L, Yield: 90% of theoretical) and the DSP yield at 90%, and evaluated changing the percentage of OPEX and CAPEX attributable to DSP from 10% to 35%. Based on these analyses, and the results shown in Figure 8, I would propose the following as reasonable commercial metrics for DEM production: Rate: > 5g/L-hr, Titer: >150g/L, Fermentation Yield: >90% of theoretical, DSP Yield: >90%, with a DSP OPEX cost target <$0.28/kg (20% of total) and DSP CAPEX target <$0.51/kg (20%) of total.

**Figure 7:**
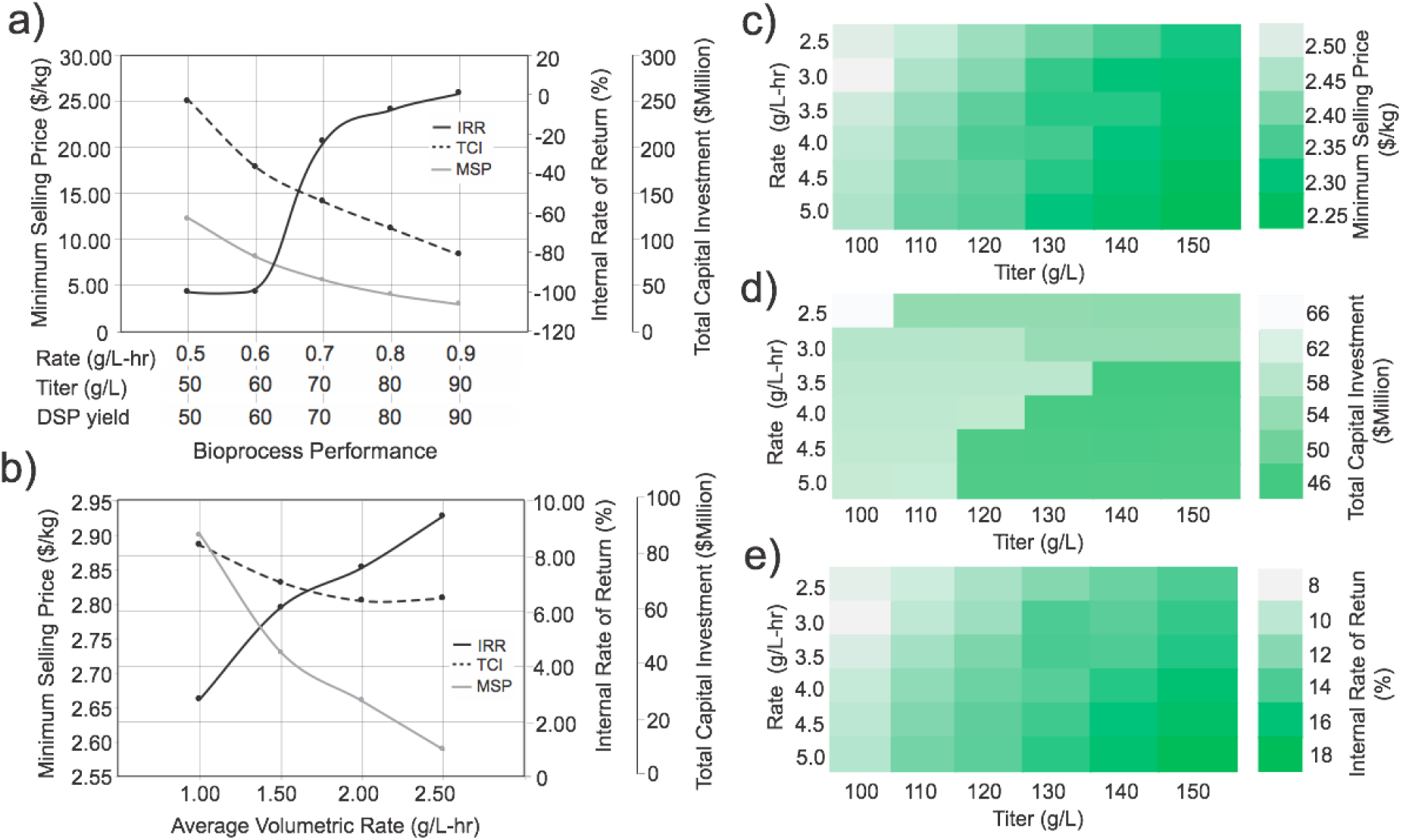
The estimated minimum selling price, total capital investment and internal rate of return for a 15kta diethyl malonate production facility as a function of bioprocess performance. Default financial and additional process assumptions were used as discussed in the main text. a) The impact of increases in average volumetric rate, titer and downstream purification yield. b) The impact of rate improvements with titer held constant at 100g/L and downstream purification yield held constant at 90%. c-e) The impact of rate, from 2.5 to 5g/L-hr, and titer, from 100-150g/L on c) minimum selling price, d) total capital investment and e) internal rate of return, respectively. In c-e) downstream purification inputs are kept constant: yield: 90%, percentage of OPEX in DSP: 15%, percentage of CAPEX in DSP: 20%.

**Figure 8:**
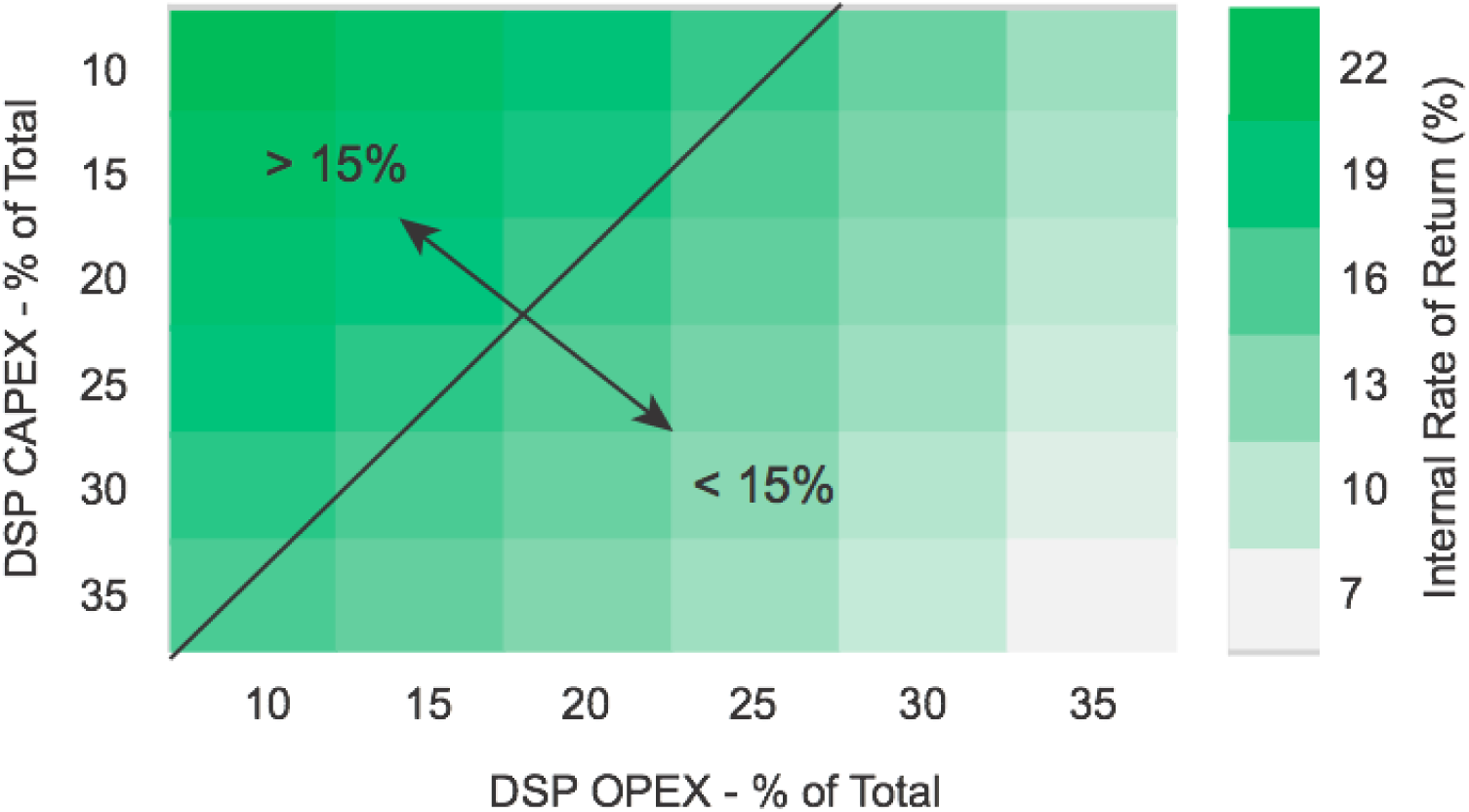
The estimated internal rate of return of a diethyl malonate production facility (15kta) as a function of the OPEX and CAPEX of downstream purification. DSP OPEX and CAPEX are both given as the percentage of the total. Default financial process assumptions were used as discussed in the main text. Fermentation performance parameters were held constant: average volumetric rate: 5g/L-hr, titer: 150 g/L, yield: 90% of theoretical. The black line indicates the division between processes that are estimated to achieve a target IRR > 15%.

## 5. Brief Conclusions and Future Directions

The Bioprocess TEA Calculator provides a generalized approach and framework to relate bioprocess performance to commercial potential. While in many aspects this is a conceptual process estimate, preliminary material and energy balances are performed for the upstream process which are used for OPEX and CAPEX calculations. These details support improved accuracy compared to an “order of magnitude” estimate. Based on my prior experience and similar approaches this conceptual cost estimation would be expected to have an accuracy +/- 30%, with respect to the upstream process.^25,26,33^ However, the impact of uncertainty in a DSP can greatly impact predictive accuracy of this tool. Missing process steps or equipment is the most common source of error in early-stage cost models. This tool is most appropriate for defining technology targets in early stage strain or bioprocess engineering programs (such as demonstrated in the case of DEM), and evaluating the relative impact of key process variables and other inputs on potential commercial success. While DEM, with a selling price <$3.00/kg, could be considered more of a bulk chemical, the calculator is also applicable to a much wider range of products most with higher values and lower production volumes.

As mentioned above, moving forward, future improvements may include a model of product acidity (or basicity), additional models of a fermentation (for example to include 2-stage production ^10^), or enabling a comparison between aerobic and anaerobic fermentations. Additionally, more specific models of several primary DSP unit operations may enable an earlier stage (conceptual) comparison of alternative purification approaches and how they may impact costs.

Even in its current form, the Bioprocess TEA Calculator enables any user to perform a basic TEA and consider commercially relevant metrics early in the development of new strains and bioprocesses. The identification of the most important variables impacting potential commercial success, even in an unoptimized analysis, can help researchers focus on the most critical research efforts. These are not always intuitive. This can be particularly true when comparing new potential bioprocesses with existing technologies. While an MSP may seem attractive, a new process must compete with an incumbent technology not only on operating costs, but on a return in investment basis, which can often represent a higher hurdle and translate to higher required performance metrics.

## Acknowledgements

I would like to acknowledge the following support: National Science Foundation (NSF): EAGER: 1445726, the Defense Advanced Research Projects Agency (DARPA): HR0011-14-C-0075, the Office of Naval Research: (ONR YIP #N00014-16-1-2558), the U.S. Dep. of Energy, Office of Energy Efficiency and Renewable Energy: (DOE EERE grant #EE0007563) and support from DMC Biotechnologies, Inc. Additionally, I would like to acknowledge input and support from the team at DMC Biotechnologies, Inc. including Matthew Lipscomb and Phil Wagner.

