## Supplementary Materials for "The Bioprocess TEA Calculator: An online techno-economic analysis tool to evaluate the commercial competitiveness of potential bioprocesses"

### Supplemental Materials

#### Section 0: Key Definitions

**OPEX:** OPEX or operating expenses, are ongoing costs for producing a given chemical and include both variable costs such as raw materials or utilities, and fixed costs such as labor. This calculator returns the operating costs per kilogram of final product produced. Complete breakdowns of OPEX can be downloaded after saving a given analysis.

**CAPEX:** CAPEX or capital expenses, are initial (as well as ongoing) expenses required for plant/facility engineering, construction and commissioning. Initial CAPEX for construction is given in $/kg or annual production capacity as well as an estimated initial total capital investment (TCI). A breakdown of capital costs can be downloaded after saving a given analysis.

**TCI - Total Capital Investment:** The TCI is the total initial cost to construct and commission the facility including both the fixed capital investment and working capital.

**MSP - Minimum Selling Price:** The minimum selling price, generated using this calculator is the selling price required to achieve the target margin upon the completion of plant ramp up (in the first year the facility is producing at nameplate production capacity). The MSP calculation includes the target margin, operating expenses, ongoing capital expenditures as well as loan payments (both principal and interest).

**Margin:** The margin input by the user is used to calculate the MSP.

**DCF - Discounted Cash Flow:** In finance, discounted cash flow analysis is a method of valuing a project using the concepts of the time value of money. Discounted cash flow analysis is widely used in investment and patent valuation. Specifically discounted cash flow is a project valuation method used to estimate the value of an investment based on its projected future cash flows. DCF analysis attempts to assess the value of an investment today, based on projections of how much money it will generate in the future.

**Selling Price:** The user input selling price, is the price used to calculate the financial returns, and should reflect the market price (or lower) for a given chemical. If the production costs (based on additional inputs) are greater than the selling price, the calculator will return negative financial results. The selling price should not be confused with the minimum selling price (MSP) which is the price for which the product would need to be sold to achieve the input margin. An MSP larger than the target selling price indicates a commercially infeasible bioprocess.

**Discount Rate**: In this case, the discount rate refers to the interest rate used in discounted cash flow (DCF) analysis to determine the net present value of future projected cash flows.

**NPV - Net Present Value:** Net present value (NPV) is the difference between the present value of projected cash inflows and the present value of projected cash outflows over a period of time. NPV is used to analyze the profitability of a projected project. NPV depends on the discount rate and accounts for the time value of money.

**IRR - Internal Rate of Return:** IRR is the annual rate of growth an investment is expected to generate. IRR is calculated using the same concept as NPV, except it sets the NPV equal to zero. IRR is used to compare potential rates of annual return for several projects over time.

**Payback Period:** In this calculator, the payback period refers to the amount of time used to perform the DCF analysis and calculate the financial returns including NPV, IRR and ROI . Simply put, the payback period is the length of time the project is given to make a return. By default for large capital projects the default payback period is 20 years.

**ROI - Return on Investment:** ROI is a performance measure used to evaluate the efficiency of an investment. To calculate ROI, the benefit (or total return) of an investment is divided by the cost of the investment. The result is expressed as a percentage or a ratio.

**Percent Debt Financed:** This input allows the user to define how much of the plant is financed (paid for with loans). The default value is 60%.

**Loan Term:** The term of the loan used to pay for plant construction

**Loan Interest Rate:** The interest rate of the loan used to pay for plant construction

**Tax Rate:** The US corporate tax rate. The United States imposes a tax on the profits of US resident corporations at a rate of 21 percent (reduced from 35 percent by the 2017 Tax Cuts and Jobs Act).

**ProForma:** A proforma (pro forma in this context means projected) business plan includes financial statements showing projected sales, operating expenses, capital costs interest and taxes.

**Net Cash Flows:** Net cash flow refers to the difference between a company's cash inflows and outflows in a given period.

**Cumulative Cash Flow:** For a given investment the cumulative cash flow simply equals the total sum of the Net Cash flow over a larger period of time, usually through the entire payback period.

**Plant/Facility Terms:**

**Capacity (kta - kilotonnes per annum):** In this calculator, the production capacity for a given plant estimate is given in kta or kilotonnes per annum. 1 kta is equivalent to 1,000,000 kg of product produced per year. While the user can input a target annual production capacity the calculator will estimate costs based on a discrete number of tanks giving an optimal capacity

**Annual Uptime:** There are 365 days (8760 hours) in a year and some of this time must be spent performing routine plant maintenance and repairs. The annual uptime is the fraction of the year where the plant is up and running. The default for the calculator is 90% or 7884 hours per year.

**Batches On Spec:** While in an ideal scenario, 100% of the product produced in a given facility is sold, in reality for most bioprocesses, issues arise with a particular batch of product wherein it does not meet the customer quality specifications (purity for example). The calculator assumes these "off-spec" batches cannot be sold and require disposal. The Batches on Spec (specification) is the percentage of the batches produced that are sold. The default for this calculator is 95%.

**Main Fermenter Size:** This calculator assumes aerobic fermentation and gives the user the choice of three large scale aerobic vessels with a total size of 250,000 Liters, 500,000 Liters and 1,000,000 Liters.

**Overall Yield:** The overall process yield is the fermentation yield multiplied by the overall yield achieved in downstream purification.

**CEPCI:** THE CHEMICAL ENGINEERING PLANT COST INDEX: The CEPCI is an index used in estimating costs in (bio)chemical process engineering. This calculator uses the CEPCI (which in 2020 is ~ 603) to estimate costs for air, water and other utilities as described by (Vasudevan and Ulrich 2006)

**TIC -** Total Installed Costs: An equipment factored capital cost estimate can be produced by taking the estimated cost of individual types of process equipment, and multiplying it by an "installation factor" to arrive at the total installed costs (TIC). In practice, this has proven to be quite a useful method since a substantial part of total project costs are made up of equipment. The installation factor includes subcontracted costs, associated direct labor costs and materials needed for installation of equipment. The TIC is equal to the equipment costs multiplied by the installation factor.

**Raw Materials:**

**Glucose Costs:** Glucose is the carbon source used in this calculator, and assumed to be DE95 dextrose (corn syrup) at a concentration of ~ 650 grams of glucose per liter. The default pricing for this feedstock as of 2020, is $0.18/lb. While many "sugar" prices available online may be anywhere from $0.25-$0.50/lb. In our experience $0.18/lb represents a reasonable starting cost from several suppliers of corn syrup. Costs as low as $0.14/lb may be achievable with longer term contracts and "over the fence" pricing. "Over the fence" pricing could be available if the new plant is co-located (next to) an existing sugar mill or other sugar production site.

**Ammonia Costs:** Ammonia is used to supply nitrogen for cell growth and or product biosynthesis (in the case where the product) contains nitrogen. Additionally, ammonia is used as the base titrant during the fermentations. Default ammonia pricing is $0.12/lb.

**Natural Gas:** Natural gas is used to generate steam and hot water for several steps in the bioprocess including sterilization and the assumed biomass heat kill decontamination step. Default natural gas pricing is $3.11/MMBtu.

**Electricity Costs:** Electricity is used to run much of the process including agitators and mixers, air blowers transfer pumps and the centrifuges used for primary cell removal.

**Chemical Inputs & Outputs**

**Theoretical Yield:** The theoretical yield is the maximal yield of a given chemical product from glucose in grams/gram. This calculation is based on the potential theoretical chemical conversion and does not assume a particular biosynthetic pathway or account for thermodynamics. Theoretical yields using a particular biochemical pathway may have a theoretical pathway yield lower than the theoretical chemical yield.

**Yield Coefficients:** Yield coefficients are calculated based upon the stoichiometry, and include those for glucose, oxygen and ammonia. For example: grams of product / gram of ammonia.

**Fermentation Inputs & Outputs**

**Rate:** The rate is the average volumetric production rate in the main fermentations. The input units are grams of product per liter per hour (g/l-hr).

**Titer:** Titer is the final concentration of the product at the end of the fermentation in grams per liter (g/L)

**Fermentation Yield:** The fermentation yield is the overall yield of product from glucose as a raw material. The calculator takes user input which is a target percentage of the theoretical yield. This is capped at 98% of the theoretical yield of the product from glucose. In any fermentation some number of cells are required to produce the product and these cells require glucose (as well as other nutrients) for replication and metabolism.

**gCDW/L:** grams of cell dry weight per liter of fermentation or cell culture broth. The calculator will estimate final biomass levels in the fermentation.

**Specific Rate:** The specific rate is the average rate of production of product per amount of biomass at the end of the fermentation and is given in units of grams of product per gCDW per hour or g/gCDW-hr. This is a metric of how good the cell line or strain is. The higher the specific rate the more the volumetric rate for a given biomass level and the higher the yield. Yields are increased because less biomass (which requires sugar) is needed to produce the same amount of product.

**Media Costs:** This calculator bases media costs on the final biomass levels that are estimated. As a result the use inputs the cost of media components per kg or biomass (kgCDW). The default value of ~$0.40/kg assumes a minimal mineral salts media, based on the FGM10 and FGM25 minimal media as reported by Menacho-Melgar et al. (Menacho-Melgar et al. 2020)

**Turnaround Time:** The turn around time is the time required to clean and ready a bioreactor for another batch, once the preceding batch is completed. This calculator defaults to a 16 hour turnaround time. This time includes the time to harvest the broth, clean the tank and lines, which is assumed to be CIP (clean in place) in this model, and refill with fresh media prior to the next inoculation.

**Oxygen Transfer Rate (OTR):** The OTR (given in mmoles of O2 per liter per hour) required in aerobic fermentations is the oxygen delivery rate needed to meet the demands of cellular respiration and product biosynthesis. Specifically the maximal OTR must match or exceed the OUR (oxygen uptake rate) of the culture. The maximal OTR as well as average OTR are calculated based on the balanced stoichiometry, as well as assumptions of rate titer and yield.. While this calculator will estimate the maximal needed OTR for a given process, it should be noted that values that exceed 250 mmol/L-hr may not be achievable in practice in large scale industrial fermentations.

**Mass Transfer Coefficient (Kla):** The mass transfer coefficient is a function of the specific bioreactor and is a measure of how well a given bioreactor can transfer oxygen. The OTR is equal to the Kla multiplied by the oxygen concentration gradient in the culture. The estimated Kla can be used to ensure that the equipment can meet the target requirements.

**Cooling Demand:** In cell culture, particularly aerobic cultures, heat generation can be significant and bioreactors must also be able to cool the culture fast enough to maintain target temperatures. The cooling demand also dictates the requirements of large scale bioreactors. Historically, the rate of heat generation in a given aerobic culture is well correlated to the oxygen consumption rate. This correlation (0.460kJ/mmole $O_2$) is used in this calculator. For more details refer to Bioprocess Engineering Principles by Doran.

**Downstream Purification Terms:**

**Primary Cell Removal:** Separation of the cells in the whole fermentation broth, to produce a "clarified fermentation broth" without cells. This calculator assumes centrifugation for initial primary cell removal and estimates the related costs (operating and capital costs).

**DSP-** Downstream Purification, including separation and purification of the target product from clarified fermentation broth.

**DSR-** Downstream Recovery, another term often used for DSP.

**Capital Terminology**:

**Equipment Costs:** The equipment costs, as the name suggests, are the costs to purchase key process equipment. Equipment costs are estimated from quotes and scaling factors. Scaling factors adjust cost estimates for the size of the equipment needed compared to that quoted. In some cases estimates are based on personal prior experiences. For details on specific sources and scaling factors, refer to the Tutorial on Capital Cost Estimation.

**Total Installed Costs (TIC):** Total Installed Costs for a piece of equipment includes an installation factor, which accounts for the estimated costs to install and commission a piece of equipment. TIC = Equipment Cost X Installation Factor. For details on specific installation factors refer to the Tutorial on Capital Cost Estimation.

**Direct Costs (DC) :** The total of direct costs include equipment and installation (TIC) additional piping and the installation of process control systems, as well as costs for site development (clearing, concrete, etc.), and the construction of warehouses and administrative buildings. The estimates include the materials and labor associated with plant construction.

**Indirect Costs :** The total of indirect costs include field expenses, project contingency, start up costs (licenses and fees), additional proratable expenses and "Home Office" charges which include engineering and project management.

**Fixed Capital Investment (FCI) :** The FCI is the sum of the Direct and Indirect Costs.

**Working Capital (WC) :** The WC is estimated at 5% of the FCI.

**Total Capital Investments (TCI) :** The TCI, which is the sum of the FCI (which is the sum of the direct and indirect costs) and the WC, is the estimated total cost of new plant construction.

### Section 1: Balancing Stoichiometry & Calculating Yields

First, we need to view the problem from the perspective of basic chemical building blocks used for biosynthesis namely $CO_2$ and $H_2$. Glucose can be thought of as a combination of these two building blocks, and the decomposition of glucose into these units is described by Equation S1.1 below. For any given organic product either carbon ($CO_2$) or reducing equivalents ($H_2$) will be the limiting reagent ( unless the product has the same balance as glucose).

Equation S1.1: $C_6H_{12}O_6 + 6\ H_2O \rightarrow 6\ CO_2 + 12\ H_2$

To balance equations for the production of a generic product from glucose as a feedstock, we can next consider each product as an assembly of building blocks, $CO_2$, $H_2$ and $NH_3$ as illustrated by Equation S1.2. We then can perform material balances to generate formulas for these coefficients.

Equation S1.2: Decomposition (a,b,c, and d are known):

$$C_aH_bO_cN_d + X\,H_2O \rightarrow Y\,CO_2 + Z\,H_2 + d\,NH_3$$

Equation S1.3: N balance: $d=d$

Equation S1.4: C balance: $Y=a$

Equation S1.5: O balance:

$X = 2Y-c$

$X = 2a-c$

Equation 6: H balance:

$b+2X \rightarrow 2Z + 3d$

$2Z = b+2X-3d$

$2Z = b+2(2a-c)-3d$

$2Z = b+4a-2c-3d$

$Z = 0.5b+2a-1c-1.5d$

In the case of glucose the $H_2$:$CO_2$ ratio is 2. For any given product this can be calculated as discussed in Equation S1.7:

Equation S1.7: $H_2$:$CO_2$ ratio = $Z/Y = 0.5(b/a)-1(c/a)-1.5(d/a)+2$

For the subset of products allowed by the calculator, there are three cases.

1) If a product has a $H_2$:$CO_2$ ratio = 2 , (neutral relative to glucose) the balanced equation is as follows

Equation S1.8: $1\ C_6H_{12}O_6 + X\ NH_3 \rightarrow Y\ C_aH_bO_cN_d + Z\ H_2O$

Equation S1.9: C balance:

$6 = aY,$

$Y=6/a,$

Equation S1.10: N balance:

$X=dY,$

$X=6(d/a)$

Equation S1.11: O balance:

$6 = cY +Z,$

$6 = 6(c/a) +Z$

$Z =6-6(c/a)$

2) If a product is more reduced than glucose (a $H_2$:$CO_2$ ratio > 2), CO2 is a needed byproduct and O2 is not a reactant. And the equation to balance takes the form of Equation S1.12.

Equation S1.13: $1\ C_6H_{12}O_6 + W\ NH_3 \rightarrow X\ C_aH_bO_cN_d + Y\ H_2O + Z\ CO_2$

Equation S1.14: N balance:

$W = dX$

Equation S1.15: H balance:

$12 + 3W = bX + 2Y$

$12 + 3dX = bX + 2Y$

$2Y = 12 + 3dX - bX$

$Y = 6 + 1.5dX - 0.5bX$

Equation S1.16: O balance:

$6 = cX + Y + 2Z$

$0 = cX + 1.5dX - 0.5bX + 2Z$

$0.5bX - cX - 1.5dX = 2Z$

$Z = 0.25bX - 0.5cX - 0.75dX$

Equation S1.17: C balance:

$6 = aX + Z$

$6 = aX + 0.25bX - 0.5cX - 0.75dX$

$6 = X(a + 0.25b - 0.5c - 0.75d)$

$X = 6/(a + 0.25b - 0.5c - 0.75d)$

3) If a product is more oxidized than glucose, O2 is a needed byproduct and O2 is not a reactant. ( a $H_2$:$CO_2$ ratio < 2), O2 is a needed reactant and CO2 is not a product. In this case the equation to balance is Equation S1.18.

Equation S1.18: $1\ C_6H_{12}O_6 + W\ NH_3 + X\ O_2 \rightarrow Y\ C_aH_bO_cN_d + Z\ H_2O$

Equation S1.19: C balance:

$6 = aY$

$Y = 6/a$

Equation S1.20: N balance:

$W = dY,$

$W = 6d/a$

Equation S1.21: H balance:

$12 + 3W = bY+2Z,$

$12 + 18d/a = 6b/a + 2Z,$

$2Z = 12 + 18d/a - 6b/a,$

$Z = 6 + 9d/a - 3b/a$

Equation S1.22: O balance:

$6 + 2X = cY + Z$

$6 + 2X = 6c/a + 6 + 9d/a - 3b/a,$

$2X = 6c/a + 9d/a - 3b/a,$

$X = 3c/a + 4.5d/a - 1.5b/a$

### Section 2: Fermentation Model

For simplicity, the calculator assumes growth associated production and a logistic growth curve. The theoretical yield (calculated as described above, Methods Section 1), as well as the user defined target percentage of this theoretical yield and the user defined titer are used to calculate the maximal biomass levels within the fermentation. If the overall fermentation yield is 90% of theoretical and achieves a given titer this dictates the available sugar (not used for product synthesis) which can be used for biomass growth, according to equations S2.1 through S2.6. In these calculations the MW of "biomass, ($C_{3.85}H_{6.69}O_{1.78}N$) " is assumed to be 95.37 g/mole, which is produced from glucose according to Equations S2.7 and S2.8, where biomass biosynthesis is assumed to achieve 80% of theoretical yield. (Battley 1987; Grosz and Stephanopoulos 1983)

Equation S2.1: Product = final Titer (g/L) * working volume (L)

Equation S2.2: Sugar to Product = Product (g/L) /theoretical Yield (g product/g glucose)

Equation S2.3: Total Sugar = Sugar to Product/(fraction Of theoretical Yield)

Equation S2.4: Sugar to Biomass (g) = Total Sugar (g) - Sugar to Product (g);

Equation S2.5: Final Biomass (gCDW) = Sugar to Biomass (g)*biomassYieldCoefficient (gCDW/g)

Equation S2.6: Final Biomass Levels (gCDW/L) =Final Biomass (gCDW) / working volume (L)

Equation S2.7: 0.84 Glucose + 1 $NH_3$ + 1.212 $O_2$ --> 1 bacteria ($C_{3.85}H_{6.69}O_{1.78}N$)+ 3.212 $H_2O$ + 1.212 $CO_2$

Equation S2.8: biomassYieldCoefficient = 0.8*(95.37)/(0.84*180.156) = 0.50 gCDW/g glucose.

The total fermentation time is calculated from the user defined final titer and average volumetric production rate (Equation S2.9). The starting biomass level is assumed to be 1% of the final biomass, based on a 1% inoculum of a similarly dense seed culture. Biomass growth is assumed to be logistic growth, where the logistic growth rate is calculated according to Equations S2.10 and S2.11. Finally the product to cell ratio, a measure of growth associated product biosynthesis, is calculated per Equation S2.12. This ratio is in effect the gram of product made per gram of biomass made, as the cell grows. This can be used to estimate a logistic product formation rate, as defined by Equation S2.13.

Equation S2.9: fermentation time (hrs) = final Titer (g/L) / ave. vol rate (g/L-hr)

Equation S2.10: A = (Final Biomass - Starting Biomass) /Starting Biomass

Equation S2.11: Logistic Growth Rate = -(log(0.01/A))/(fermentation time), (hr-1)

Equation S2.12: Product to cell ratio = Titer /(finalBiomass-startingBiomass)

Equation S2.13: Logistic Production Rate = Product to cell ratio * Logistic Growth Rate

The above equations can then be used to estimate key fermentation variables, such as biomass levels, and product concentration as a function of time. This fermentation time course can be used to estimate additional operating expenses which vary as a function of the fermentation (See Section 3 below). It is also worth noting that in the case of logistic growth the maximal rate of increase in biomass (and product) occurs when the biomass is equal to ½ of the final biomass. This maximal point is used to estimate process maximums such as oxygen transfer and cooling demands.

**Section 3: Operating Cost Estimates**

Plant operating costs include variable operating costs, such as raw materials and utilities as well as fixed operating costs including labor and additional fixed costs, each will be discussed in turn below. Some additional assumptions used for several of the calculations include a main fermentation working volume ratio of 0.85, and a fermentor aspect ratio of 3.0.

*Raw Materials*

The calculator estimates the raw materials needed given the target plant capacity and fermentation performance as well as other user defined inputs. These include the costs of water, glucose, ammonia, and the costs of CIP (clean in place reagents). Glucose and ammonia costs are estimated as described in the main text. The costs of media components are estimated based on the final biomass levels in the main production fermentations (kgCDW). The default value of ~$0.40/kgCDW assumes a minimal mineral salts media, based on the FGM25 minimal media (which can support biomass levels of 25gCDW/L) as reported by Menacho-Melgar et al and detailed below in Table S3.1. (Menacho‑Melgar et al. 2020)

| Table S3.1 Media Costs for FGM25 Media (Menacho‑Melgar et al. 2020) | | | | | |
|---|---|---|---|---|---|
| Component | Concentration ($kg/m^3$) | Unit Price ($/kg) | Source | Cost ($/1000L of media) | Cost ($/kgCDW) |
| Iron(II) Sulfate | 0.0243056 | 1.21 | https://www.icis.com/chemicals/channel-info-chemicals-a-z/ | 0.029 | 0.000 |
| Magnesium Sulfate | 0.300915 | 0.40 | https://www.icis.com/chemicals/channel-info-chemicals-a-z/ | 0.119 | 0.005 |
| Calcium sulfate | 0.0085088 | 0.008 | http://www.spectrumanalytic.co | 0.000 | 0.000 |

| | | | | | |
|---|---|---|---|---|---|
| | | | m/support/library/rf/Gypsum.htm | | |
| Thiamine | 0.01 | 70.00 | https://dir.indiamart.com/impcat/thiamine-hydrochloride.html | 0.700 | 0.028 |
| Ammonium sulfate | 22.5 | 0.168 | https://www.icis.com/chemicals/channel-info-chemicals-a-z/ | 3.771 | 0.151 |
| Trace Metals | Not a significant cost at levels used. | | | | |
| Citric acid | 1 | 1.65 | https://www.icis.com/chemicals/channel-info-chemicals-a-z/ | 1.654 | 0.066 |
| monopotassium phosphate | 0.6845126 | 1.54 | Alibaba | 1.057 | 0.042 |
| dipotassium phosphate | 0.865774 | 1.54 | Alibaba | 1.336 | 0.053 |
| **Total** | | | | $8.666 | $0.346 |

The costs for the clean in place (CIP) of the main fermentation vessels is based on two cleaning solutions:  a caustic solution (NaOH) used at a final concentration of 2 weight % and a peracetic acid sterilizing solution used as a working concentration of 0.02%. The calculator assumes the tanks are cleaned after each batch with a full volume of both cleaning solutions in turn. Bulk costs for NaOH are estimated at $0.15/kg ($150/tonne). Costs for peracetic acid are estimated at $5/L of 20% peracetic acid, which is diluted to a working concentration of 0.02%.

*Fermentation Utilities*

The Bioprocess TEA Calculator estimates the required utilities needed for the bioprocess including water, compressed air and costs of mass transfer, cooling water, as well as electricity and heat for key unit operations. Cost estimates for the majority of the utilities in this calculator are based on utility estimations by Ulrich & Vasudevan (Vasudevan and Ulrich 2006). The estimates rely on the Chemical Engineering Plant Cost Index (CEPCI a metric of inflation, https://www.chemengonline.com/pci-home) , and the "Cost of Fuel, ($/GJ)" which is in turn estimated based on the user defined natural gas cost according to Equation S3.1. Additionally, the calculator has default inputs for natural gas ($3.10/MMBtu), electricity ($0.11/Kwh) and CEPCI (603) based on current estimates.

Equation S3.1: CostOfFuel = NaturalGasCost/1.05505 ($/GJ)

The process water requirements are estimated based on the average water demand rate as dictated by the media requirements of the main fermentation as defined below in Equations S3.2 through S3.4. Water costs are estimated per Ulrich & Vasudevan (Vasudevan and Ulrich 2006), based on the needed flow rates and the CEPCI and Cost of Fuel (Equation S3.3).

Equation S3.2: Demand Rate = ((annual media Volume/annual Fermentation Up Time) *m3/sec*

Equation S3.3: Water Costs =(0.0007+0.00003*(Demand Rate^-0.6))*CEPCI + 0.02*CostOfFuel (*$/m3)*

Equation S3.4: Annual Water Costs = annual media Volume*waterCost, *$*

As the calculator assumes an aerobic fermentation, the costs of air and air delivery (mass transfer) are critical. These costs are estimated as follows. Firstly, similarly to the cases of glucose and ammonia, as described above, the total oxygen required for both biomass and product synthesis (in the case where a product is more oxidized than glucose and requires molecular oxygen for synthesis) are estimated based on the stoichiometry. In the case of biomass growth, the model assumes biomass is synthesized at 80% of theoretical yield, the additional glucose consumed (lost to $CO_2$) is assumed to be completely oxidized to carbon dioxide at the expense of molecular oxygen (Equation S3.5). This in turn is used to estimate the average needed airflow rate assuming 9.375 moles of pure $O_2$ per L of air (Equation S3.6). We can then estimate the average airflow requirement per Equation S3.7. In this calculation we assume that only 75% of the oxygen can be consumed in the fermentation (i.e the dissolved oxygen is set to a 25% set point and the tanks are not run under conditions of zero residual oxygen). (Benz 2008) The cost of compressed air per $m^3$ is again estimated per Ulrich & Vasudevan (Vasudevan and Ulrich 2006), in Equation S3.8, where the maximum fermenter pressure is calculated from the height of the tanks, based on an aspect ratio of 3 and the user defined tank volume. This leads to a final estimate of compressed air costs (Equation S3.9).

Equation S3.5: Cumulative $O_2$ = $O_2$ needed for biomass + $O_2$ needed for product + $O_2$ lost to $CO_2$

Equation S3.6: Cumulative Air = Cumulative $O_2$/(9.375*1000) ($m^3$)

Equation S3.7: ave. Annual Airflow = (Cumulative Air/(Annual Uptime (hrs)*3600))*1.333, ($m^3$/s)

Equation S3.8: Air Cost per $m^3$ =

(0.00005*(ave. Annual Airflow^-0.3))*log(max Ferm. Pressure)*CEPCI) + 0.0009*log(max Ferm. Pressure)*Cost Of Fuel *($/m3)*

Equation S3.9: Annual Cost Of CompressedAir = Air Cost per $m^3$*Cumulative Air, ($/year)

The above estimates and calculations also enable us to estimate an average and maximal needed oxygen transfer rate as well as the costs of mass transfer according to Equations S3.10 through S3.19. Given logistic growth (and knowing maximal uptake occurs at ½ of the target biomass level) the maximal OTR attributable to biomass growth can be calculated per Equations S3.11 and S3.12, and the stoichiometry given above in Equation S2.7. A similar calculation is performed for product synthesis, again only in the case where $O_2$ is needed for product synthesis, and the maximal OTRs required for biomass and product synthesis are added. The maximal required oxygen flow rate (and subsequently the maximal air flow rate) can be estimated from the max OTR and the vessel working volume (Equations S3.13 & S3.14). In Equation S3.14, again we account for the fact that practically only a maximum of 75% of the oxygen in the air may be consumed. The maximum required mass transfer coefficient ($k_{LA}$) for the bioreactors is estimated in Equations S3.15 and S3.16. This again assumes a maximal driving force for oxygen delivery wherein the residual $O_2$ is solution is kept at 25% of saturation, wherein saturation is estimated at ~ 0.2 mmoles of $O_2$ per liter of media.

Equation S3.10: ave. OTR = (cum. $O_2$/ (annual Up Time * Total Annual WorkingVolume) (mmole/L-hr)

Equation S3.11: biomassYieldCoefficient$O_2$ = (95.37)/(1.212*32), (g biomass/g $O_2$)

Equation S3.12: $MaxOTR_{Biomass}$=

(((1/biomassYieldCoefficient$O_2$)*(1000/32)*(Logistic Growth Rate)/4)*(final Biomass), (mmoles $O_2$/ L-hr)

Equation S3.13: Max Oxygen Flow Rate = maxOTR* Vessel Working Volume, (mmoles/hr per tank)

Equation S3.14: Max AirFlow Rate = max Oxygen Flow Rate/9.375/0.75/1000/3600, ($m^3$/sec)

Equation S3.15: max $O_2$ gradient = 0.2mmoles/L - 0.05mmoles/L = 0.15 mmoles/L

Equation S3.16: maxKla = (maxOTR/0.15)/3600, (sec-1)

Finally we can estimate the costs of the required mass transfer. This is done based on estimations of mass transfer costs in aerobic bioreactors by Humbird, Davis and McMillan (Humbird, Davis, and McMillan 2017). Briefly, the cost of mass transfer is estimated as a function of the power required for mass transfer, which in turn is estimated as a function of the mass (kg) transferred. For stir tank reactors we estimate a power requirement of 1.8 kW/kg $O_2$ 1.8 kW/kg $O_2$. The mass fraction of $O_2$ in air is 0.233. The cumulative power required for air transfer is then multiplied by electricity costs to estimate the total mass transfer costs.

Equation S3.17: Cumulative Air = cumulativeAir*1.225, (conversion to kg) , from Equation S3.6.

Equation S3.18: Annual Mass Transfer Power = 1.8*0.233*(cumulative Air), (kWh) (Humbird, Davis, and McMillan 2017)

Equation S3.19: Annual Cost of Mass Transfer = annual Mass Transfer Power*Electricity Cost, ($)

Next we can estimate the costs of bioreactor cooling. In general the cooling requirement in aerobic fermentations is well correlated with oxygen consumption. (Doran 1995) Specifically we can estimate the heat generated (cooling needed) in the fermenter per Equation S3.20. Given the cooling requirement we can then estimate the associated costs again per Ulrich & Vasudevan (Vasudevan and Ulrich 2006), in Equations S3.24 through S3.26. Cooling water is assumed to be 29℃ (~85℉). (LePree 2010; Buecker 2017) If the fermentation temperature is greater than 33℃, the return temperature is assumed to be 4℃ lower than the fermentation temperature. If the fermentation temperature is lower than 33℃, the return temperature is assumed to be 30℃, 1 degree higher than the cooling water temperature. The difference between the cooling water temperature and the return temperature is “dT” in equation S3.24. Based on the annual demand and average flow rate the cost of cooling water is again estimated per Ulrich & Vasudevan (Vasudevan and Ulrich 2006), Equations S3.26.

Equation S3.20: Heat generation = 0.460 (kJ/ mmole $O_2$ consumed)

Equation S3.21: Cumulative Cooling Demand = 0.460*cumulative $O_2$ (kJ)

Equation S3.22: max Cooling Rate = 0.460*maxOTR (kJ/L-hr)

Equation S3.23: ave. Cooling Rate = Cumulative Cooling Demand /Up Time (kJ/hr)

Equation S3.24: Cooling Water = Cum. Cooling Demand /(4.184*(dT))/1000, (m3)

Equation S3.25: ave. Cooling Water Flow Rate = (Cooling Water /Up Time)/3600, (m3/sec)

Equation S3.26: annual Cost Of Cooling Water =

Cooling Water *((0.0001+(0.00003/ave. Cooling Water Flow Rate))*CEPCI + 0.003*CostOfFuel), ($)

We can also estimate the costs of media sterilization and the heat kill of used biomass at the end of the fermentation. These estimates are treated in Equations S3.27 and S3.28. In these calculations we assume a sterilization temperature of 120 ℃, and an ambient temperature of 25 ℃, as well as a sterilization efficiency of 20%. Similarly, we can estimate the cost of heat treating the waste biomass (Equations S3.29 through S3.30). In this case we assume a heat kill temperature of 60 ℃. To estimate the volume of concentrated cells that must be heat treated, we need to account for the fact that in order to be able to discharge solids they must be fluid. This means that as cells are discharged from disc stack centrifuges (which as discussed below are assumed in this process) they take some broth/water with them. In general for fermentation broth, centrifuge solids can be assumed to carry 35-40% water with them. So for every 10gCDW separated, we assume 6 g or 6mL of water. The total water % by weight = 6/(16) = 37.5% (includes volume of cells). As a result we can estimate the volume that must be heat treated as a function of the final biomass concentration per Equation S3.31.

Equation S3.27: Media Sterilization Energy Consumption = 4.184*mediaVolume*0.2*(120-25) , (kJ/year)

Equation S3.28: Cost of Sterilization = (Media Steri. Energy/1055056)*NaturalGasCost, ($/year)

Equation S3.29: Heat Kill Energy Consumption = 4.184*heatKillVolume*0.2*(60-25) ,

Equation S3.30: Cost of Heat Kill = (Heat Kill Energy /1055056)*NaturalGasCost, ($/year)

Equation S3.31: heatKillVolume = 1.6*Final Biomass (gCDW)/1000, (Liters)

*Primary Cell Removal Utilities*

As we have above for the fermentation process area, we can also estimate the cost of utilities for primary cell removal. This involves estimating the costs associated with centrifugation, which as mentioned above are assumed to be disc stack centrifuges. The centrifuge flow rate for an individual centrifuge is assumed to be 10,000 L/hour (in an acceptable range for disc stack centrifuges, (Szepessy and Thorwid 2018; Doran 1995)). The total annual volume to be centrifuged in the total fermentation volume to be processed is estimated per the annual fermentation volume. The needed hours of centrifugation can be calculated based on our flow rate, Equation S3.32, which we can then use to calculate the needed number of centrifuges and the actual estimated centrifuge up time (Equations S3.33 and S3.34). We can then estimate the power consumption of a disc stack centrifuge operation at these flow rates per Szepessy &

Thorwid, (Szepessy and Thorwid 2018) which then can be used to estimate the utility costs of centrifugation (Equations S3.35 through S3.37).

Equation S3.32: hours of Centrifugation required = annual Centrifuge Volume/10000, (hours)

Equation S3.33: number Centrifuges = Ceiling(hours of Centrifugation /annualUpTime)

Equation S3.34: Actual Centrifuge Uptime = annual Centrifuge Volume/(number Centrifuges*10000),

Equation S3.35: Power Cons. per centrifuge = 0.3(kW/$m^3$/hr)*(10$m^3$/hr)=3.0, (kW)

Equation S3.36: Total Centrifuge Power Consumption = 3.0* number Centrifuges, (kW)

Equation S3.37: Cost of Centrifugation = Total Cent. Power *Actual Uptime*ElectricityCost, ($)

*Downstream Purification Operating Costs.*

Lastly we can turn to the estimated operating costs associated with downstream purification. Operating costs for DSP are considered as a fraction of the total operating costs per Equations S3.38 and S3.39, where "Fermentation Opex" in this case includes primary cell removal. The user defines the fraction of the total percentage of OPEX due to DSP. While this is dependent on the specifics of the process and DSP, as mentioned above we recommend using a range from 10% at the low end to 40% at the high end. The default values for DSP of 20% which is recommended when the user does not have a specific downstream in mind.

Equation S3.38: Total Operating Costs = Fermentation Opex/(1-dspOPEXfraction);

Equation S3.39: DSP Opex = dspOPEXfraction*Total Operating Costs

Labor and other fixed are estimated according to David et al (R. E. Davis et al. 2018) and described in the main text.

### Section 4: Capital Cost Estimates

As described in the main text, based on the fermentation model and other process inputs as well as estimated operating expenses calculated above, the size of major required equipment can be estimated. Specifically, the size and or number of tanks, transfer pumps, heat exchangers and other key items including centrifuges, tanks and pumps for water treatment and processing, boilers, cooling towers (and cooling tower pumps), and air handling (compressors and receivers) are all estimated. Once needed equipment sizes are estimated, a standard approach to capital estimation is used, which relies on scaling factors and installation factors (as well as accounting for inflation) according to Equations 4.1 through 4.3 , details of these factors are given in Table S4.1 below.

Equation S4.1: $Purchase\ Cost\ =\ Quoted\ Cost\ *\ (Actual\ Size/Quoted\ Size)^{Scaling\ Factor}$

Equation S4.2: $Total\ Installed\ Cost\ =\ Inflation\ Factor\ *\ (Purchase\ Cost\ )\ *\ Installation\ Factor$

Equation S4.3: $Piping\ Cost\ =\ 0.045\ *\ Total\ Installed\ Cost$

| **Table S4.1: Equipment Cost Estimates** | | | | | | | |
|---|---|---|---|---|---|---|---|
| **Equipment** | **Quoted Cost** | **Actual/ Quoted Size** | **Quote Year** | **Scaling Factor** | **Inf. Factor** | **Inst. Factor** | **Source** |
| **Main Fermentation Costs** | | | | | | | |

| | | | | | | | |
|---|---|---|---|---|---|---|---|
| Main Fermenters | $176,000 | 1 | 2009 | 0.7 | 1.13 | 2 | (R. E. Davis et al. 2018) |
| Agitators | $36,000 | Variable | 2013 | 0.5 | 1 | 1.5 | |
| Main Transfer Pump | $3,900 | Variable | 2009 | 0.8 | 1.17 | 2.3 | |
| Feed Storage Tank | $70,000 | Variable | 2009 | 0.7 | 1.17 | 2.6 | |
| Feed Transfer Pump | $3,900 | Variable | 2009 | 0.8 | 1.17 | 2.3 | |
| Base Storage Tank (Ammonia) | $98,000 | Variable | 2010 | 0.7 | 1.13 | 1.5 | |
| Base Transfer Pump (Ammonia) | $3,900 | Variable | 2009 | 0.8 | 1.17 | 2.3 | |
| Acid Storage Tank (sulfuric) | $196,000 | Variable | 2010 | 0.7 | 1.13 | 2.0 | |
| Acid Transfer Pump | $3,900 | Variable | 2009 | 0.8 | 1.17 | 2.3 | |
| Dry Chemical Addition | $100,000 | 1 | 2020 | 1 | 1 | 2 | This study |
| Agitated Media Prep | $91,200 | Variable | 2009 | 0.7 | 1.17 | 2.6 | (R. E. Davis et al. 2018) |
| MediaTransfer Pump | $3,900 | Variable | 2009 | 0.8 | 1.17 | 2.3 | |
| CIP Tanks (3) | $98,000 | Variable | 2010 | 0.7 | 1.13 | 2.0 | |
| CIP Transfer Pump (3) | $3,900 | Variable | 2009 | 0.8 | 1.17 | 2.3 | This study |
| Main Ferm. Piping | 0.045 * Total Installed Cost of Main Ferm Equipment | | | | | | |
| **Seed Fermentation Costs** | | | | | | | |

| | | | | | | | |
|---|---|---|---|---|---|---|---|
| Seed Train | 0.27 * Total of Installed Costs for Main Fermentation | | | | | | (R. Davis et al. 2013) |
| **Primary Cell Removal** | | | | | | | |
| Centrifuges | $325,000 | 1 | 1998 | NA | 1.59 | 1.8 | |
| Broth Storage Tank | $1,317.000 | Variable | 2010 | 0.7 | 1.0 | 1.8 | (R. E. Davis et al. 2018) |
| Transfer Pumps | $3,900 | Variable | 2009 | 0.8 | 1.17 | 2.3 | |
| Cell Removal Piping | 0.045 * Total Installed Cost of Primary Cell Removal Equipment | | | | | | |
| **Utilities** | | | | | | | |
| Cooling Tower | $1,3750,00 | Variable | 2010 | 0.6 | 1.12 | 1.5 | (R. E. Davis et al. 2018) |
| Cooling Tower Pumps | $283,671 | Variable | 2010 | 0.8 | 1.12 | 3.1 | |
| Boiler Pack (250psig) | $100,000 | Variable | 1998 | 0.6 | 1.59 | 2.0 | (Loh et al. 2002). |
| Air Compressor (50psig) | $1318600 | variable | 2014 | 1.00 | 1.03 | 1.6 | (R. E. Davis et al. 2018) |
| Air Receiver (50psig) | $17,000 | variable | 2010 | 0.7 | 1.12 | 3.1 | |
| Air Dryer | $15,000 | variable | 2009 | 0.6 | 1.17 | 1.8 | |
| Process/Municipal Water Tank | $250,000 | variable | 2009 | 0.6 | 1.17 | 1.7 | |
| Water Softener System | $78,000 | variable | 2009 | 0.6 | 1.17 | 1.8 | |
| Water Pumps | $15,292 | variable | 2009 | 0.6 | 1.17 | 3.1 | |

| | | | | | | | |
|---|---|---|---|---|---|---|---|
| Wastewater Storage Tank | $1,317.000 | Variable | 2010 | 0.6 | 1.0 | 1.8 | |
| Wastewater Pump | $3,900 | Variable | 2009 | 0.6 | 1.17 | 2.3 | |
| Potable Water System | $75,000 | 1 | 2020 | NA | 1 | 1.7 | This study |
| Heat Exchangers | $15,000 | Variable | 1998 | 0.5 | 1.59 | 3.1 | (Loh et al. 2002) |
| Utilities Piping | 0.045 * Total Installed Cost of Process Utility Equipment | | | | | | |
| **Control Systems** | | | | | | | |
| Electrical and Control systems estimated at 10% of total installed costs. | | | | | | | This study |

*Main Fermentation Capital Estimates*

The costs of the main aerobic fermenters were scaled based on estimates from prior estimates. (R. Davis et al. 2013)These tank cost estimates include cooling coil costs, but agitator costs were estimated according to tank main vessel size. These agitators are sized to handle any reasonable mass transfer rate. One transfer pump (exit) is assumed per fermentation tank. The addition tanks (glucose, acid and base) tanks were sized to hold up to 12 hour of feed/additions based on calculations discussed above (Section 3) Each addition tank has an associated transfer pump scaled appropriately. Additionally a single agitated media prep tank, the same size of the main fermenters is shared among the production fermenters.  Lastly 3 CIP tanks were sized to 1/100th of the main fermentation vessel to hold concentrated cleaning solutions. Associated transfer pumps as well as a heater and filter for CIP are also included.

*Seed Fermentation  Capital Estimates*

The capital costs of the seed train was estimated at 27% of the total installed costs of the main fermentation area, excluding CIP associated costs.(R. Davis et al. 2013) The seed fermentations are assumed to be cleaned by SIP (sterilization in place) , requiring a steam supply.

*Primary Cell Removal & Broth Storage Capital Estimates*

Disc stack centrifuges with a flow rate of 10,000L/hr were used for cost estimates. 10,000 L/hr translates to 10$m^3$/hr or 0.0025 $m^3$/s. These centrifuges were designed to completely harvest bacterial cells with a sedimentation velocity $u_o$*=6.81x10-9 m/s.* (Doran 1995) With these assumptions, the required centrifuge sigma factor can be estimated per Equation S4.4,

Equation S4.4: $= Q/2uo = 205580\ m^2/s$

where Q is the flow rate. The purchase cost of a disc stack centrifuge, with a sigma of 200,000 $m^2$/s is estimated at $325,000 in 1998, or accounting for inflation, $516,750 in 2020. (Petrides et al. 2019; Autor Harrison et al. 2003) The number of needed centrifuges is estimated based upon the annual fermentation volume. Installation factor for these centrifuges is estimated at 1.8. A broth storage tank was sized to be large enough to hold 50% of the total main fermentation volume.

*Process Utility Capital Estimates*

Process utility requirements are estimated as described in the estimation of operating costs, discussed in Section 2 above. The associated capital estimates are discussed here. There are five primary process utilities considered : 1) cooling water, 2) air handling, 3) steam generation (ie a boiler), 4) water supply and 5) wastewater pretreatment. Capital costs for these utility systems are estimated as discussed in turn below.

*Cooling* - The annual demand for cooling (average kJ/hr) is calculated based on the fermentation calculations. A cooling tower is used to provide cooling water at 29℃. The flow rate of cooling water needed is estimated as discussed in Section 3 above. The capital costs are estimated based on quotes and scaling factors (scaled to water demand) as annotated in Table S4.1. The pumps required for cooling water recirculation are similarly estimated based on previously reported estimates. (R. E. Davis et al. 2018) Piping again is estimated at 4.5% of the installed costs of the pump and tower.

*Steam generation & Heat Exchangers*- Steam is used in two primary places in the current bioprocess design (not accounting for steam use in any downstream processing), including 1) the sterilization of the media, lines and seed (SIP) fermenters and 2) the heat treatment of the waste biomass. The costs and energy needed in these steps is estimated as detailed in Section 3. These estimates can enable us to estimate the total amount of steam needed for these heaters and or heat exchangers. In both cases a batch heating model is used to estimate steam requirements. (Baldwin et al. 1995) A package boiler producing steam at 250 psig (he ~1910kJ/kg) was assumed, and active sterilization and heat kill were assumed to account for 5% of the annual plant uptime. The estimated cost of the industrial boiler package was scaled based on capacity per Table S4.1 and cost estimates from Loh et al. (Loh et al. 2002). The sizes of two plate heat exchangers were estimated for media sterilization and heat treatment respectively according to Equations S4.13 through S4.15, with a conservative overall heat transfer coefficient U = 1 $kW/m^2$-K, and where ldT is the log mean temperature difference. The cost of the exchangers was scaled based on area per Table S4.1.

Sterilization

Equation S4.5: Time = (0.05)* Annual Fermentation Uptime, (hrs)

Equation S4.6: mass = Annual Volume of Ferm. broth(L)*(1.05kg/L), (kg)

Equation S4.7: Heat transfer rate required = 4.19*(120-25)*mass/time

Equation S4.8: $Steam_{STERILIZATION}$ = 2.2046*Heat transfer rate/1910, (lb/hr)

Heat Kill

Equation S4.9: mass = Heat Kill Volume (L)*(1.1kg/L), (kg)

Equation S4.10: Heat transfer rate required = 4.19*(60-25)*mass/time

Equation S4.11: $Steam_{HEAT\ KILL}$ =2.2046*Heat transfer rate/1910, (lb/hr)

Totals

Equation S4.12: Total Steam = $Steam_{STERILIZATION}$ +$Steam_{HEAT\ KILL}$

Equation S4.13: ldT = ((201-Temp)-(Temp-25))/(Math.log((201-Temp)/(Temp-25)))

Equation S4.14: ExchangerArea = Heat transfer rate/(U*ldT);

*Air Handling* - The air handling consists of three primary pieces of equipment 1) an air dryer, 2) an air compressor and 3) the air receiver for distribution of compressed air to process equipment, which are the fermentation vessels in this case. The air receiver was designed to hold air at a maximum pressure of 50 psig and a minimum pressure 20% higher than the maximal fermentation pressure. The maximal fermentation pressure is based on the height of the vessels based on an aspect ratio of 3 and the user input vessel volume and 50 psig is high enough for even the largest tank in this calculator. The receiver was then sized according to Equation S4.13 where the hold time ( for the receiver to go from the maximal to minimum pressure was set to 30 seconds). The cost of the air compressor is estimated based on required compressor power per Douglas et al. (Douglas and Douglas 1988; Luyben 2018) The needed compressor power is estimated according to Equations S4.14 and S4.15, assuming an inlet at 25℃ and 1 atmosphere and an outlet at the bottom of the main fermenters. Air is assumed as the only gas with an isentropic coefficient for air (y) = 1.4 and a molecular weight of 38.96. Additionally, the compressor was assumed to have an efficiency of 70% (0.7). Lastly the instrument air dryer was scaled based flow rate using previously reported cost estimates. (R. E. Davis et al. 2018)

Equation S4.13: Volume = (hold time) *Ave. Annual Airflow*14.7/(50-1.5*Max Ferm pressure)

Equation S4.14: Power = (2.31*(y/(y-1))*0.7/28.96)*(comp Outlet T-comp Inlet T)*airflow

Equation S4.15: comp Outlet T = comp Inlet T*(50/1.5*Max Ferm pressure)^(y(y-1)/y))

*Water and Wastewater* - Water supply was assumed to be from a municipal source, with costs assumed for a holding tank, water softener system (as well as associated pumps) and a potable water system estimated per Table S4.1. Similarly, costs for a wastewater holding tank and transfer pumps were estimated again per Table S4.1.

**Section 5: Financial Calculations**

With estimations of OPEX and CAPEX as discussed above financial calculations can be performed. These are focused on generating annual cash flows based on costs and potential revenues. To begin, there are some assumptions that are used, which are currently not editable by the user. These include assumptions related to plant construction and start up as well as depreciation, loan payments and the accounting basis. Construction is assumed to take 2 years from the start of the project to complete, with 70% of the capital spent in the first year and 30% in the second. After year 2, ongoing capital expenses (for repairs and maintenance) are assumed at 10% of the initial capital investment. After construction, production is assumed to take 3 years to ramp up, with 50% of nameplate capacity produced in the first year of production, and 75% of nameplate capacity produced in the second year of production, reaching 100% of nameplate capacity produced in the third year of production. Depreciation is assumed to be a 10 year straight line depreciation. Loan payments on the plant financing are interest only during the construction period. Lastly, EBITDA, EBIT, EBT and net cash flows are calculated using standard formulas. The MSP is calculated so that a 30% margin is achieved after the ramp up period is completed. NPV, ROI and IRR are all calculated using standard approaches.

Battley, Edwin H. 1987. *Energetics of Microbial Growth*. Wiley.

Benz, Gregory T. 2008. “Piloting Bioreactors for Agitation Scale-Up.” *Cephalalgia: An International Journal of Headache*. http://cmbe.engr.uga.edu/bche3180/Benz%202008%20Chem%20Eng%20Prog.pdf.

Buecker, B. 2017. “Cooling Tower Heat Transfer Fundamentals [WWW Document].” In *Power Eng*.

Davis, R., L. Tao, E. C. D. Tan, M. J. Biddy, G. T. Beckham, C. Scarlata, J. Jacobson, et al. 2013. “Process Design and Economics for the Conversion of Lignocellulosic Biomass to Hydrocarbons: Dilute-Acid and Enzymatic Deconstruction of Biomass to Sugars and Biological Conversion of Sugars to Hydrocarbons.” https://doi.org/10.2172/1107470.

Davis, Ryan E., Nicholas J. Grundl, Ling Tao, Mary J. Biddy, Eric C. Tan, Gregg T. Beckham, David Humbird, David N. Thompson, and Mohammad S. Roni. 2018. “Process Design and Economics for the Conversion of Lignocellulosic Biomass to Hydrocarbon Fuels and Coproducts: 2018 Biochemical Design Case Update; Biochemical Deconstruction and Conversion of Biomass to Fuels and Products via Integrated Biorefinery Pathways.” https://doi.org/10.2172/1483234.

Doran, Pauline M. 1995. *Bioprocess Engineering Principles*. Elsevier.

Douglas, James Merrill, and Merrill Douglas. 1988. *Conceptual Design of Chemical Processes*. McGraw-Hill.

Grosz, R., and G. Stephanopoulos. 1983. “Statistical Mechanical Estimation of the Free Energy of Formation of E. Coli Biomass for Use with Macroscopic Bioreactor Balances.” *Biotechnology and Bioengineering* 25 (9): 2149–63.

Humbird, D., R. Davis, and J. D. McMillan. 2017. “Aeration Costs in Stirred-Tank and Bubble Column Bioreactors.” *Biochemical Engineering Journal* 127 (November): 161–66.

LePree, Joy. 2010. “JUST COOL IT!” *Chemical Engineering* 117 (6): 17.

Loh, H. P., EG&G Technical Services, Inc., Morgantown, WV (United States), Jennifer Lyons, and Charles W. White. 2002. “Process Equipment Cost Estimation, Final Report.” https://doi.org/10.2172/797810.

Luyben, William L. 2018. “Capital Cost of Compressors for Conceptual Design.” *Chemical Engineering and Processing - Process Intensification* 126 (April): 206–9.

Menacho‐Melgar, Romel, Zhixia Ye, Eirik A. Moreb, Tian Yang, John P. Efromson, John S. Decker, Ruixin Wang, and Michael D. Lynch. 2020. “Scalable, Two‐stage, Autoinduction of Recombinant Protein Expression in E. Coli Utilizing Phosphate Depletion.” *Biotechnology and Bioengineering* 26 (June): 44.

Petrides, Demetri, Doug Carmichael, Charles Siletti, and Alexandros Koulouris. 2019. “Bioprocess Simulation and Economics.” *Essentials in Fermentation Technology*. https://doi.org/10.1007/978-3-030-16230-6_9.

Szepessy, Stefan, and Peter Thorwid. 2018. “Low Energy Consumption of High-Speed Centrifuges.” *Chemical Engineering & Technology* 41 (12): 2375–84.

Vasudevan, Palligarnai T., and Gael D. Ulrich. 2006. “How to Estimate Utility Costs.” *Chemical Engineering* , no. 4: 66–69.
